# Filter inference: A scalable nonlinear mixed effects inference approach for snapshot time series data

**DOI:** 10.1101/2022.11.01.514702

**Authors:** David Augustin, Ben Lambert, Ken Wang, Antje-Christine Walz, Martin Robinson, David Gavaghan

## Abstract

Variability is an intrinsic property of biological systems and is often at the heart of their complex behaviour. Examples range from cell-to-cell variability in cell signalling pathways to variability in the response to treatment across patients. A popular approach to model and understand this variability is nonlinear mixed effects (NLME) modelling. However, estimating the parameters of NLME models from measurements quickly becomes computationally expensive as the number of measured individuals grows, making NLME inference intractable for datasets with thousands of measured individuals. This shortcoming is particularly limiting for snapshot datasets, common e.g. in cell biology, where high-throughput measurement techniques provide large numbers of single cell measurements. We extend earlier work by Hasenauer et al (2011) to introduce a novel approach for the estimation of NLME model parameters from snapshot measurements, which we call filter inference. Filter inference is a new variant of approximate Bayesian computation, with dominant computational costs that do not increase with the number of measured individuals, making efficient inferences from snapshot measurements possible. Filter inference also scales well with the number of model parameters, using state-of-the-art gradient-based MCMC algorithms, such as the No-U-Turn Sampler (NUTS). We demonstrate the properties of filter inference using examples from early cancer growth modelling and from epidermal growth factor signalling pathway modelling.

**Author summary:** Nonlinear mixed effects (NLME) models are widely used to model differences between individuals in a population. In pharmacology, for example, they are used to model the treatment response variability across patients, and in cell biology they are used to model the cell-to-cell variability in cell signalling pathways. However, NLME models introduce parameters, which typically need to be estimated from data. This estimation becomes computationally intractable when the number of measured individuals – be they patients or cells – is too large. But, the more individuals are measured in a population, the better the variability can be understood. This is especially true when individuals are measured only once. Such snapshot measurements are particularly common in cell biology, where high-throughput measurement techniques provide large numbers of single cell measurements. In clinical pharmacology, datasets consisting of many snapshot measurements are less common but are easier and cheaper to obtain than detailed time series measurements across patients. Our approach can be used to estimate the parameters of NLME models from snapshot time series data with thousands of measured individuals.

## Introduction

Variability is an intrinsic property of biological systems and is often the reason for their complex behaviour [1]. Examples are plentiful. One is the evolution of organisms, whereby variability in the genetic material across individuals is one of the key drivers for adaptation of populations [2]. Another is the human adaptive immune system, wherein variability in the antigen binding sites across antibodies is crucial for the defence against a large variety of pathogens [3]. However, variability in the function and regulation of cells is also the cause of many diseases, such as cancer and Altzheimer’s disease [4–6]. Quantifying variability is therefore central to understanding many biological systems.

Nonlinear mixed effects (NLME) modelling is a popular approach to model variability in populations [7, 8]. NLME models introduce a set of model parameters which typically need to be estimated from measurements. However, the inference of NLME models from measurements quickly becomes prohibitively expensive when the number of measured individuals increases [9]. This shortcoming of NLME inference is particularly limiting when individual entities can only be measured once, since such ‘snapshot’ measurements do not capture individual trajectories and are therefore relatively uninformative about the dynamics across individuals, requiring large numbers of snapshot measurements for good inference results. Such datasets are, however, typically intractable for NLME inference. In this article, we introduce a novel inference approach, which we call filter inference. We demonstrate that filter inference provides a scalable NLME inference approach for snapshot time series measurements.

Snapshot measurements are particularly common in cell biology, where experimental techniques, such as single-cell RNA sequencing and flow cytometry, provide high-throughput measurements without the possibility to repeatedly measure individual cells [10–12]. The availability of snapshot measurements paired with the limitations of NLME inference has led to the development of a variety of inference methods. Hasenauer et al (2011) simulate measurements to construct an approximate likelihood for the model parameters using kernel density estimation (KDE) [9]. To this end, they make explicit assumptions about the population parameter distribution. Dixit et al (2020) use the simulated measurements to fit the histograms of the observed measurements exactly, making no explicit assumptions about the population parameter distribution [13]. Instead, they require that the entropy of the model parameter distribution is maximised. Lambert et al (2021) use exhaustive simulations from a prior distribution of the model parameters to construct a contour volume distribution which enables an efficient inference of the model parameters [14]. Similar to Dixit et al’s method, Lambert et al’s approach also does not make any explicit assumptions about the population parameter distribution. However, it does require the simplifying assumption that measurement noise is negligible for the inference. Browning et al (2022) use an ABC inference approach to infer NLME models from snapshot measurements, where summary statistics of simulated and observed measurements are compared [15].

With filter inference, we extend Hasenauer et al’s inference approach. In particular, we introduce a differentiable form of the approximate likelihood, making it applicable for state-of-the-art gradient-based sampling algorithms, such as Hamiltonian Monte Carlo (HMC) and the No-U-Turn sampler (NUTS) [16–18]. This improves the inference efficiency, especially for NLME models with many parameters. In addition, we generalise the approach to an ABC framework that uses filters instead of summary statistics. This allows us to systematically study the properties of filter inference and the consequences of different filter choices on the parameter estimates.

The body of this article is divided into two sections: a methods and a results section. In the methods we review the NLME modelling framework and introduce filter inference. In the results we demonstrate the performance of filter inference for two NLME inference problems, which both suppose access to snapshot measurements: 1. for an early cancer growth model; and 2. for an epidermal growth factor (EGF) pathway model. We also use these modelling problems to demonstrate the reduction of the computational costs when using filter inference, and draw comparisons between filter inference and summary statistics-based ABC. We conclude the article by addressing potential sources for information loss and bias. The data, models and scripts used in this article are hosted on https://github.com/DavAug/filter-inference. A user-friendly API for filter inference has been implemented in the open source Python package `chi` [19].

## Methods

NLME models acount for the dynamics of heterogeneous populations using a hierarchical modelling structure [7, 8]. First, a time series model, 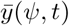, is used to model the dynamics of an individual. Here, 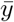 denotes a quantity of interest, *t* denotes the time and *ψ* denotes the parameters of the model. An example time series model for early cancer growth is illustrated in red in Fig 1, where the quantity of interest,

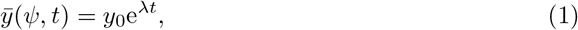

captures a patient’s tumour volume over time. The parameters of the model, *ψ* = (*y*_0_, *λ*), are the initial tumour volume and the growth rate.

**Fig 1.**
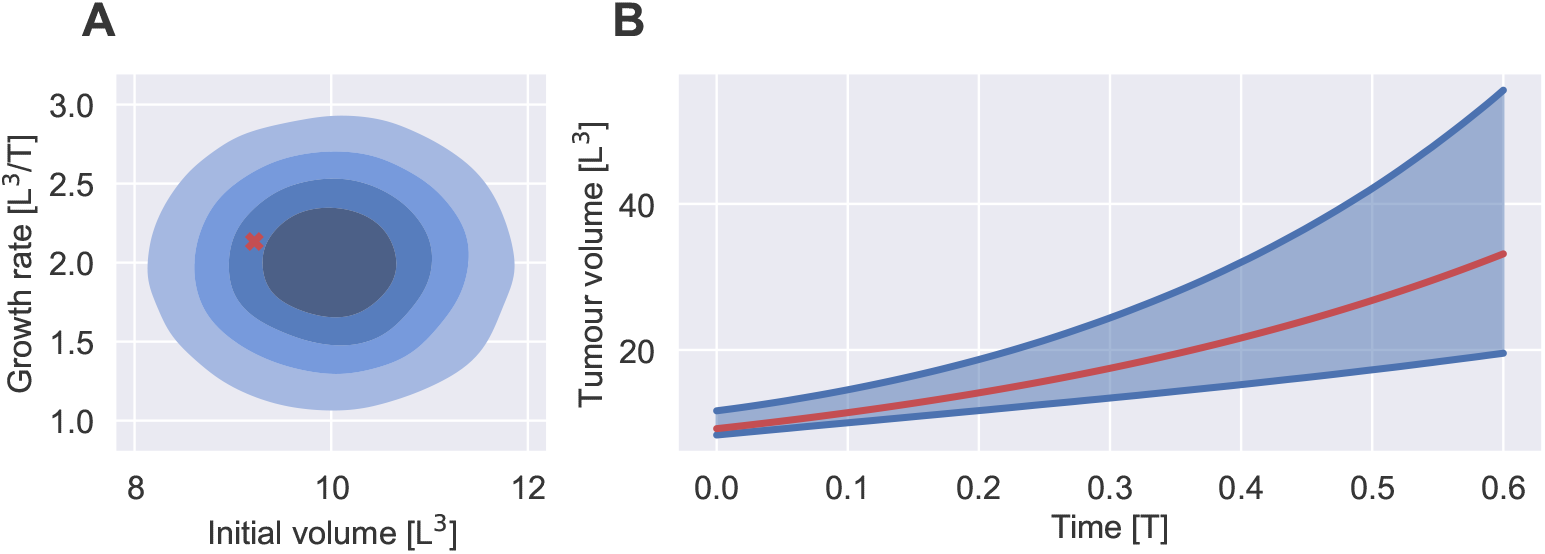
NLME model of early cancer growth. A: Shows the population model, *p*(*ψ* | *θ*), for *θ* = (10, 1, 2, 0.5) in shades of blue and the parameters of a randomly chosen individual in red. The shades of blue indicate the bulk 20 %, 40 %, 60 % and 80 % probability of the distribution. B: Shows the distribution of tumour volumes across individuals in the population, *p*(*y* | *θ, t*), in blue and the tumour volume of a randomly chosen individual over time in red. The blue lines indicate the 5th and 95th percentile of the tumour volume distribution at each time point. The quantities are shown in arbitrary units. *L* denotes length dimensions and *T* denotes time dimensions.

Second, a population model, *p*(*ψ*|*θ*), with population parameters *θ* is used to capture the inter-individual variability (IIV) by modelling the distribution of *ψ* in the population. For example in Fig 1A, the initial tumour volume and the growth rate are normally distributed across patients

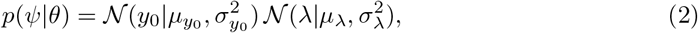

where 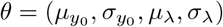 denotes the population means and standard deviations of *ψ*. For clarity, we will refer to *ψ* as individual-level parameters and to *θ* as population-level parameters. Although equivalent, note that our notation deviates from the standard NLME literature, where the individual-level parameters are decomposed into fixed and random effects, 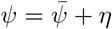 [7]. In this notation, we recover the population model in Eq 2, by letting the fixed effects, 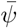, be equal to the population means, and the random effects, *η*, be normally distributed with zero means and variances equal to the population variances.

With this hierarchical structure, the variability in the dynamics can be simulated by repeatedly evaluating the time series model for different samples of the individual-level parameters. Each sample of *ψ* represents an individual in the population. As the number of samples increases, the histogram over the simulations converges to the population distribution of the quantity of interest, which is illustrated in blue in Fig 1B. Formally, this population distribution is defined as

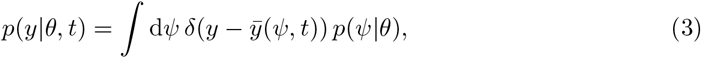

where *δ*(*x*) denotes the Dirac delta distribution. In the cancer growth example, *p*(*y*|*θ, t*) quantifies the probability with which a randomly chosen individual in the population has a tumour of volume *y* at time *t*. In general, the spread of *p*(*y*|*θ, t*) quantifies the IIV of the quantity of interest. NLME models assume that each set of individual-level parameters, *ψ*, fully characterises the dynamics of an individual. As a result, the heterogeneity of the dynamics in the population arises exclusively from the population model, *p*(*ψ*|*θ*).

### Nonlinear mixed effects inference

The NLME model, as defined in Eq 3, is fully characterised by the population parameters *θ*. For most biological modelling problems these parameters are unknown and need to be estimated from data. Many algorithms and software packages for the inference of NLME models have been developed and excellent reviews exist [19–22]. Here, we will review the Bayesian inference approach, partly to exposit existing NLME inference approaches but also to introduce concepts necessary to understand filter inference.

In order to infer parameters from measurements, it is customary to include an error model, 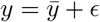, in the NLME model definition [7]. The error model accounts for both measurement noise and discrepancies between the model and the true process – these two processes are collectively accounted for by *ϵ*. This extends the deterministic time series model output to a distribution of measurements, 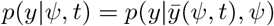. For ease of notation, we extend the definition of *ψ* to also include the parameters of the error model. A common model choice is assuming normally distributed residual errors, 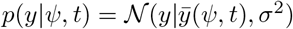. The distribution of measurements across individuals in the population can then be defined analogously to Eq 3

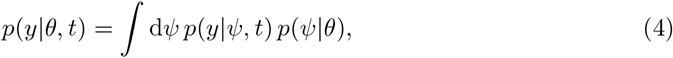

where *p*(*y*|*ψ, t*) replaces the Dirac delta distribution. Thus, Eq 3 can be interpreted as a special case of Eq 4, where measurements are assumed to capture the value of the time series model output, 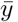, without any error. Note that Eq 4 implicitly defines a joint probability distribution for measurement and individual-level parameters, *p*(*y, ψ*|*θ, t*) = *p*(*y*|*ψ, t*) *p*(*ψ*|*θ*), which is marginalised over *ψ* on the right hand side of Eq 4: *p*(*y*|*θ, t*) = ∫ d*ψ p*(*y, ψ*|*θ, t*).

Given measurements across individuals, the joint probability distribution, *p*(*y, ψ*|*θ, t*), can be used to define a hierarchical log-likelihood for the model parameters

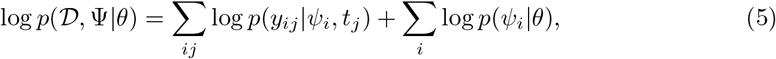

quantifying the likelihood of parameter values, (*θ*, Ψ), to capture the observed dynamics, 𝒟 = (*Y, T*). We use Ψ = (*ψ*_1_, *ψ*_2_, …, *ψ*_*N*_) to denote the individual-level parameters across *N* measured individuals, and *Y* to denote the associated matrix of measurements across individuals and times. In particular, the *ij*th element of *Y, y*_*ij*_, denotes the measurement of individual *i* at time *t*_*j*_, where the vector of all unique measurement times is denoted by *T*. As a result, *Y* has *N* rows, and *K* = dim(*T*) columns. Missing measurement values do not contribute to the likelihood. Eq 5 shows that the hierarchical likelihood comprises a term accounting for the likelihoods of individual-level parameters to describe the measurements, and a term accounting for the likelihood of the population parameters to describe the distribution of the individual-level parameters.

An example dataset suitable for the inference of the early cancer growth model is outlined in Table 1. For simplicity, we neglect challenges of the measurement process and assume that it is possible to measure the tumour volume across patients *in vivo*. In practice, it may be more feasible to use the *in vitro* proliferation of cancerous cells from tissue samples as a proxy for the tumour growth. For inference, the dataset can be expressed in matrix form

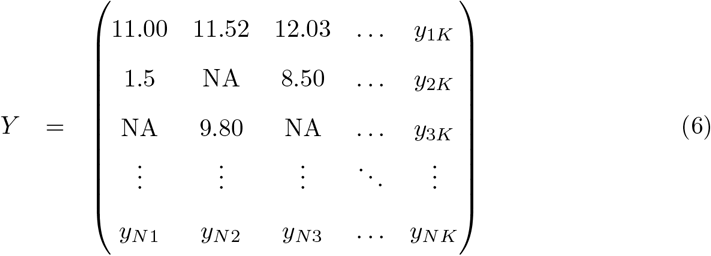

with *T* = (1.5, 2.1, 4, …, *t*_*K*_), where the first row contains the measurements of the patient with ID 1, the second row contains the measurements of the patient with ID 2, and so on. NA denotes missing values. An important feature of the dataset is that individuals are not necessarily measured with the same frequency, or at the same time points. In the extreme, the dataset may contain only one measurement per individual, i.e. snapshot measurements.

**Table 1.** Outline of an example tumour volume dataset. The dataset contains (fictitious) time series measurements of tumour volumes across patients. Patients are labelled with unique IDs. The time and tumour volume are presented in arbitrary units. *T* indicates the time dimension and *L* the length dimension.

| | ID | Time $[T]$ | Tumour volume $[L^3]$ |
| --- | --- | --- | --- |
| 1 | 1 | 1.5 | 11.00 |
| 2 | 2 | 1.5 | 8.30 |
| 3 | 1 | 2.1 | 11.52 |
| 4 | 3 | 2.1 | 9.80 |
| 5 | 1 | 4 | 12.03 |
| 6 | 2 | 4 | 8.50 |
| ⋮ | ⋮ | ⋮ | ⋮ |

In a Bayesian inference approach, Bayes’ rule is used to translate the hierarchical likelihood into a distribution of parameter values consistent with the observations and prior knowledge, also known as the posterior distribution

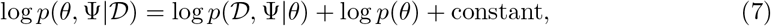

where *p*(*θ*) is the prior distribution of the population-level parameters [23]. *p*(*θ*) is used in Bayesian inference to quantify knowledge about parameter values and is a modelling choice. The model parameters can now be inferred from *p*(*θ*, Ψ| 𝒟) using sampling algorithms, such as Markov chain Monte Carlo (MCMC) algorithms [18], see Alg 1.

### Filter inference

The intractability of NLME inference for snapshot data stems from the increasing cost of evaluating the log-likelihood as the number of measured individuals grows. This is because the evaluation of the log-likelihood defined in Eq 5 requires one evaluation of the time series model for each observed individual, resulting in computational costs that increase at least linearly with the number of measured individuals. This expense renders NLME inference intractable when thousands of individuals are measured, especially when time series models are defined by systems of differential equations that need to be solved numerically.

#### Algorithm 1

NLME inference using Metropolis-Hastings MCMC sampling. The details of the proposal and acceptance step are omitted for clarity, but may be found in [24].

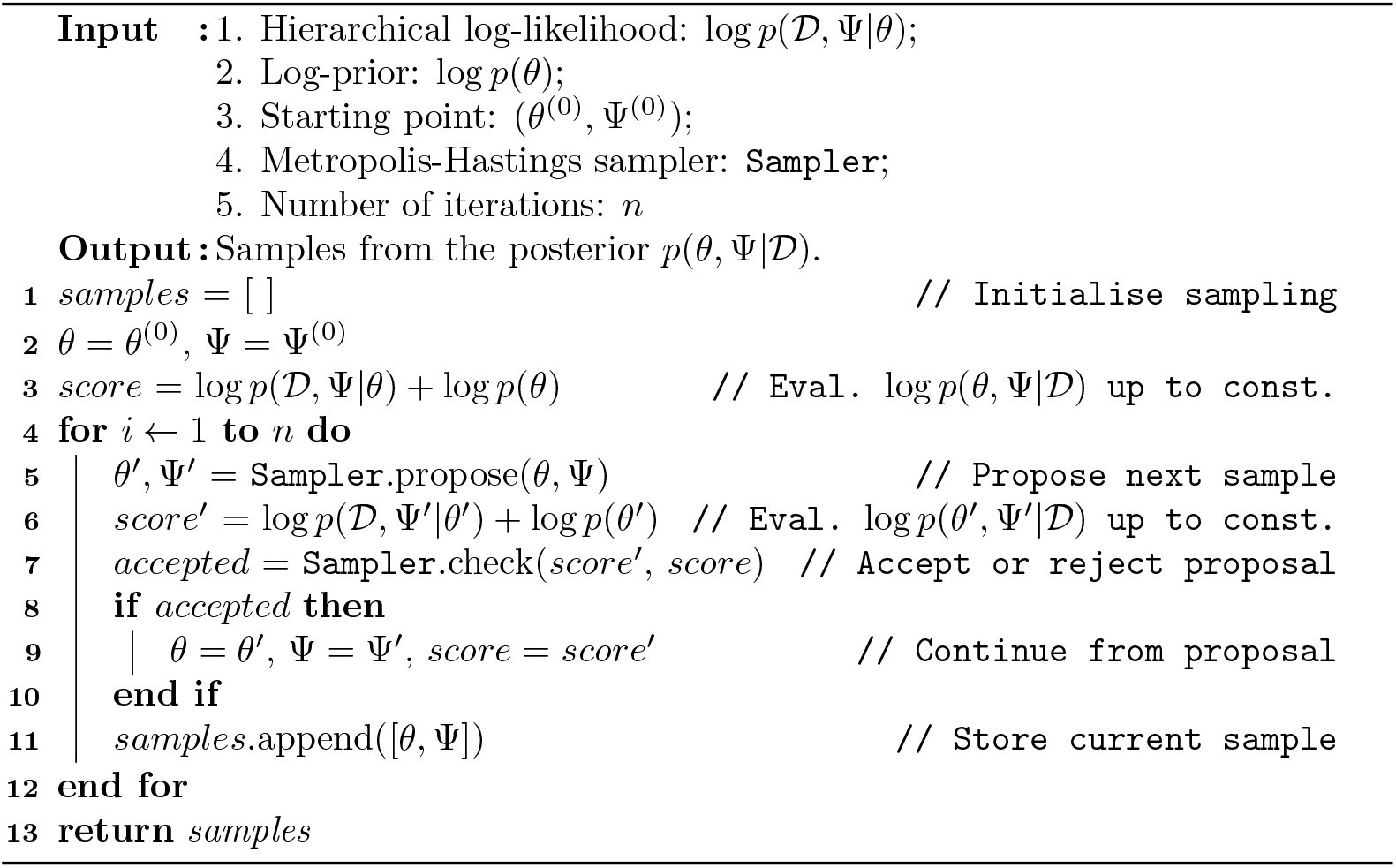

In theory, this intractability can be avoided by fitting to the measurements on a population-level, removing the need to evaluate the time series model for each individual separately. In particular, using the population distribution of measurements defined in Eq 4, a log-likelihood for the population-level parameters can be defined directly, log *p*(𝒟 |*θ*) = ∑_*ij*_ log *p*(*y*_*ij*_|*θ, t*_*j*_). From this population-level log-likelihood, we can derive a log-posterior, log *p*(*θ*| 𝒟) = log *p*(𝒟 |*θ*) + *p*(*θ*) + constant, which can be inferred using MCMC sampling. However, in practice, the integral in Eq 4 is too expensive to compute to make *p*(*y*|*θ, t*) tractable for inference.

ABC is a popular inference method for problems with intractable likelihoods [25]. Here, simulations of measurements are compared to the observed measurements to construct an approximate log-likelihood. Standard ABC methods do this by computing the distance between summary statistics of the simulated and observed measurements. The distance is quantified using kernel functions whose acceptance scale is defined by a manually chosen error margin. An alternative ABC approach, called Bayesian synthetic likelihood (BSL), uses parametrised distributions to construct synthetic likelihoods for the simulated summary statistics [26–28].

Filter inference is a novel variant of ABC that uses filters to estimate the likelihood of model parameters from the observed measurements directly. In particular, filters approximate the measurement distribution at different time points, *p*(*y*|*θ, t*), based on simulated measurements. These approximations are then used to estimate the log-likelihood

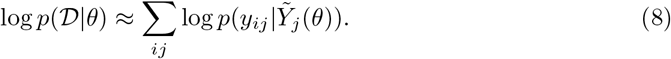

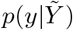 denotes the filter, and 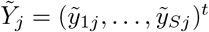 denotes *S* simulated measurements at time *t*_*j*_. Different filter choices capture different information from the real and simulated measurements. For example, a Gaussian filter, introduced below in Filters, compares the mean and variance of the measurements, much like standard ABC based on the mean and variance. However, in contrast to standard ABC, the error margin of filter inference is set automatically, see Relationship to summary statistics-based ABC. The algorithmic details of filter inference are presented in Alg 2.

Alg 2 uses MCMC sampling, similar to Alg 1, to infer the posterior distribution. The main difference between the algorithms is the replacement of the hierarchical log-likelihood evaluation by the estimate of the population-level log-likelihood, defined in Eq 8. In particular, we estimate the log-likelihood by simulating measurements from the model, *p*(*y*|*θ, t*), by first sampling simulated individuals, 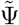, from the population model, *p*(*ψ*|*θ*), see Alg 2 lines 17-21. We then simulate measurements for each simulated individual by sampling from *p*(*y*|*ψ*_*s*_, *t*) in lines 24-29. Here, we use *s* to label simulated individuals, instead of *i* which we reserve for real individuals. Using the simulated measurements, we construct a filter that summarises population-level information of the measurements. The details of this construction are filter-specific and are discussed below. The filter defines a population-level distribution of measurements, which we use to estimate the likelihood of the model parameters, see lines 21-29. From this estimate we can derive an estimate of the posterior which is computationally tractable.

#### Algorithm 2

Filter inference using Metropolis-Hastings MCMC sampling. The details of the proposal and acceptance step are omitted for clarity, but may be found in [24].

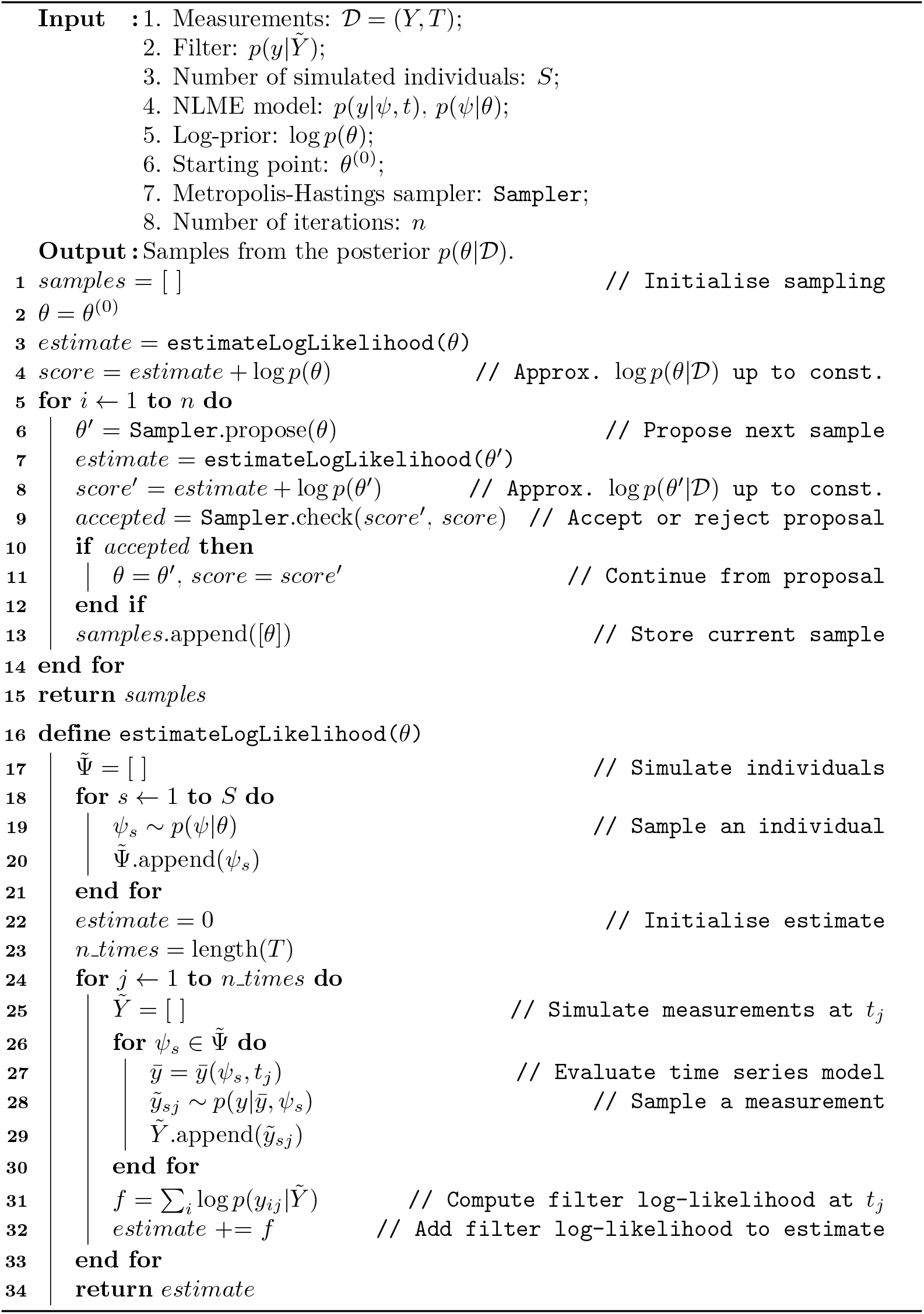

The main innovation of filter inference is to make the number of time series model evaluations independent of the number of observed individuals. In this way, filter inference remains tractable even when millions of snapshot measurements are used for parameter estimation. In particular, in Alg 2 the number of time series model evaluations is determined by the number of simulated individuals, *S*, and the number of measured time points, see line 24 and its surrounding for-loops. In an optimised implementation, this number can be reduced to a total of *S* time series model evaluations per log-likelihood estimation, see S1 Appendix. The dominant computational costs of filter inference therefore do not scale with the number of measured individuals, but instead are set by the number of simulated individuals.

This form of approximate inference was first introduced by Hasenauer et al for a specific filter choice: the lognormal KDE filter introduced below [9]. Alg 2 generalises this approach to an ABC framework, where filters can be chosen specific to the needs of the inference problem. However, Hasenauer et al’s algorithm reportedly becomes inefficient for models with more than a few parameters [13]. This is because the approach samples from the posterior using the Metropolis-Hastings (MH) MCMC algorithm, whose sampling efficiency is known to scale poorly with the dimension of the posterior distribution [24]. Our current generalisation does not resolve this limitation, as it also uses the MH algorithm for sampling.

Efficient sampling algorithms for high dimensional models exist, such as the Hamiltonian Monte Carlo (HMC) MCMC algorithm and its variants [16, 18]. HMC uses gradient-information to produce better proposals, resulting in a higher sampling efficiency per step. However, the estimate of the log-likelihood from Eq 8 is a stochastic function of the population-level parameters *θ*, and therefore is not differentiable. In particular, the estimation of the log-likelihood involves random sampling from the population model and from the individual-level measurement distributions, see lines 19 and 27 in Alg 2. As a result, estimates of the log-likelihood will vary due to the stochasticity inherent in both the population-level distribution and the measurement noise distribution, even for fixed population-level parameters. This makes the estimate non-differentiable, and thus HMC cannot be used to improve the efficiency of the algorithm.

However, we can recast the log-likelihood estimate from Eq 8 into a differentiable form by defining a joint distribution of measurements, simulated measurements and simulated individual-level parameters, 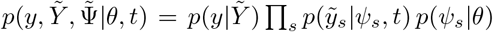. From this joint distribution, we can define a hierarchical log-likelihood comprising the filter estimate of the population-level log-likelihood, and the log-likelihood of model parameters 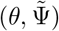 to describe the simulated measurements 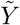

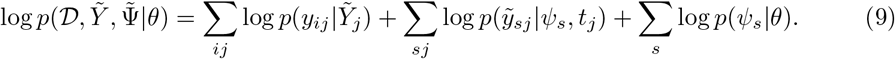

We can use this log-likelihood and Bayes’ rule to derive a log-posterior, 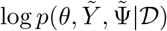, analogously to the hierarchical log-likelihood and the NLME log-posterior in Eqs 5 and 7. This log-posterior is differentiable with respect to its parameter dimensions. As a result, we can use HMC to efficiently sample from 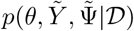 even for high-dimensional NLME models. Once 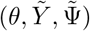 are inferred, the approximate posterior for the population-level parameters can be obtained by considering only the *θ* estimates (i.e. marginalisation). The algorithmic details of the approach are presented in Alg 3. The algorithm is illustrated using a MH sampler for easier comparison with Algs 1 and 2. Note that line 11 in Alg 3 already implements the marginalisation over 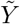 and 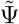.

#### Algorithm 3

Filter inference (differentiable form) using Metropolis-Hastings MCMC sampling. The details of the proposal and acceptance step are omitted for clarity, but may be found in [24].

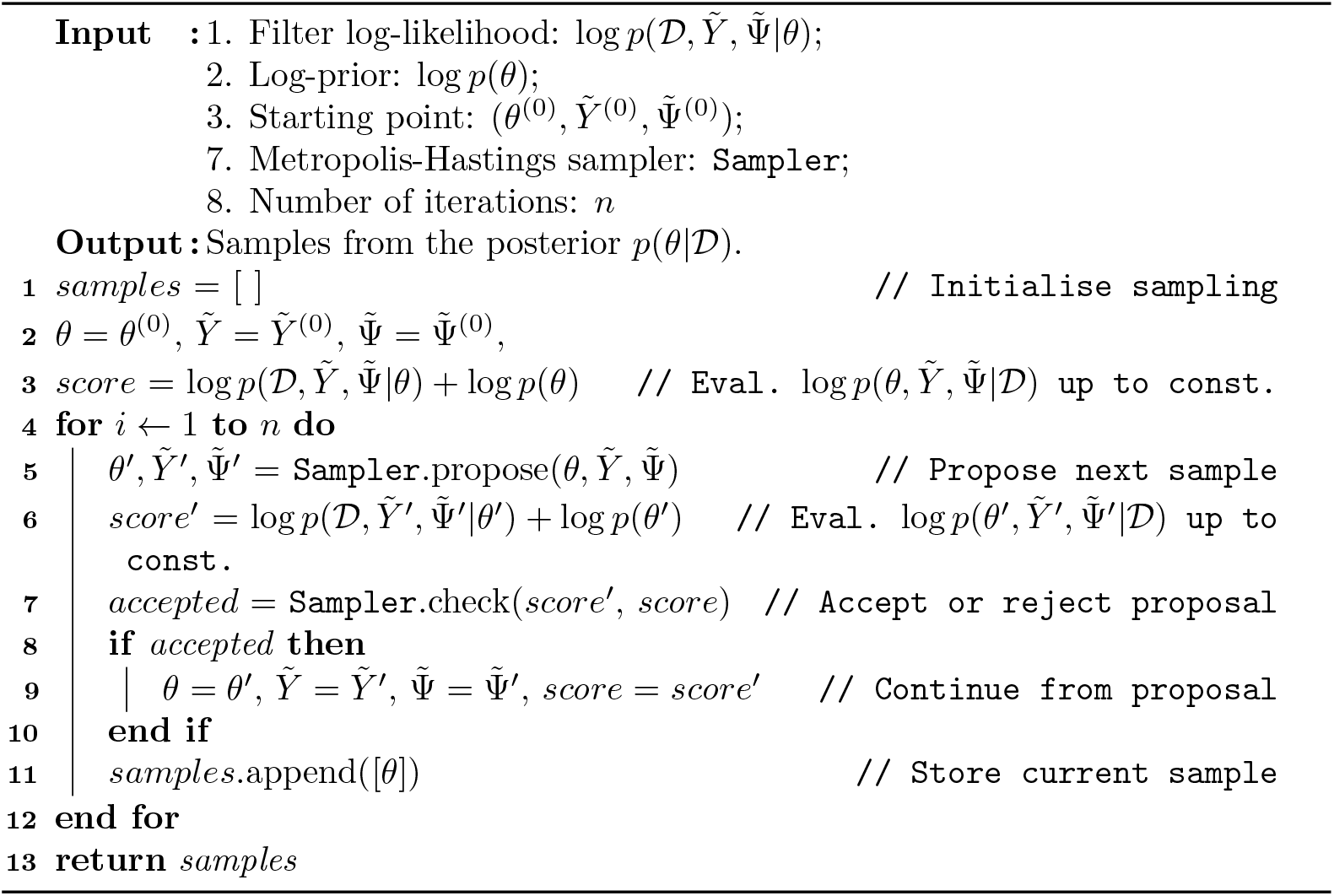

Importantly, the posteriors for *θ* inferred from the non-differentiable likelihood, Eq 8, and from the differentiable likelihood, Eq 9, are identical. The main difference is that the implementation of the non-differentiable form uses ancestral sampling to simulate individuals and measurements to estimate the log-likelihood, which makes the estimate a non-differentiable, stochastic function of the population-level parameters. This non-differentiability is eliminated in Eq 9 by explicitly formulating a log-likelihood term for each random variable contributing to the overall likelihood of the population-level parameters. In this way, the likelihood becomes a deterministic function of the random variables 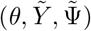. This completes the definition of filter inference.

### Filters

Filters are the central element of filter inference, making inference of NLME models from measurements of thousands of individuals possible. Below we introduce five filters that we have found useful in our experiments.

#### 1. Gaussian filter

A Gaussian filter summarises the population measurement distribution using a Gaussian distribution

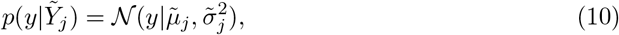

where 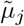 and 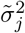 are given by the empirical mean and variance of the simulated measurements, 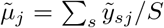 and 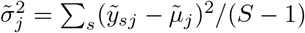, where *S* denotes the number of simulated individuals.

A Gaussian filter is illustrated with other filters in Fig 2, where we simulate *S* = 100 measurements from the early cancer growth model at *t* = 1, using the population model introduced in Early cancer growth model inference. We use the simulated measurements to construct the filters, e.g. for the Gaussian filter we compute the mean and variance of the simulations. A random realisation of each filter is illustrated in red. Repeating this construction 1000 times, we estimate the 5th to 95th percentile of the filter density distribution, illustrated in blue. As a reference for the filter approximations, the figure shows the exact population measurement distribution, *p*(*y*|*θ, t*), in black.

**Fig 2.**
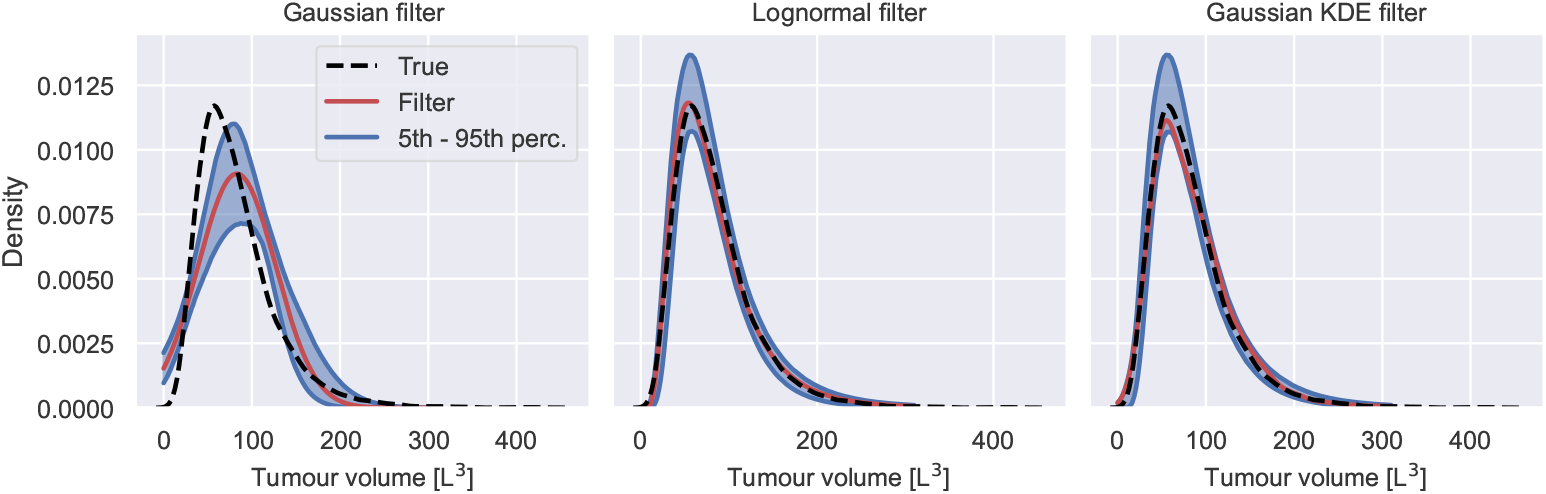
Filters in filter inference. The figure shows a Gaussian filter, a lognormal filter and a Gaussian KDE filter of the early cancer growth model for *S* = 100 simulated individuals at time *t* = 1. Each filter is illustrated by a randomly chosen realisation, illustrated in red, and the 5th to 95th percentile of the filter distribution for different sets of simulated individuals. As a reference, the exact population measurement distribution is illustrated in black.

#### 2. Lognormal filter

A lognormal filter summarises the population measurement distribution using a lognormal distribution

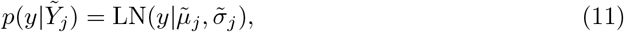

where the location and scale of the lognormal distribution, 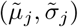, are given by the empirical mean and standard deviation of the log-transformed simulated measurements.

#### 3. Gaussian mixture filter

A Gaussian mixture filter summarises the population measurement distribution using a Gaussian mixture distribution

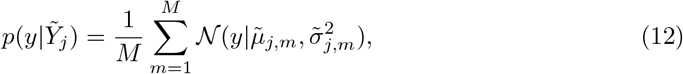

where *M* is a hyperparameter and determines the number of Gaussian kernels. The mean and variance of the Gaussians are estimated from the simulated measurements by assigning the simulated individuals uniformly to *M* subpopulations. The empirical means and variances are estimated from each subpopulation separately.

#### 4. Gaussian KDE filter

A Gaussian KDE filter summarises the population measurement distribution using a Gaussian kernel density estimation

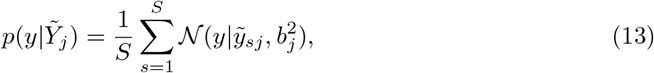

where *S* is the number of simulated individuals. A Gaussian KDE population filter is a Gaussian mixture population filter, where each individual is assigned to its own subpopulation, i.e. *M* = *S*. The bandwidth of the kernels, 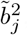, is a hyperparameter of the population filter. In this article we use the widely used rule of thumb for bandwidth selection, 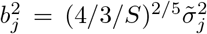, following Hasenauer et al (2011) [9]. Here, 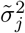 is the empirical variance of the simulated measurements at time *t*_*j*_.

#### 5. Lognormal KDE filter

A lognormal KDE filter summarises the population measurement distribution using a lognormal kernel density estimation

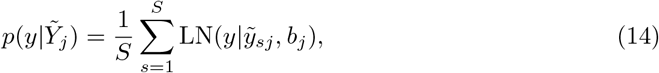

The bandwidth is computed using the rule of thumb, 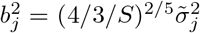, where 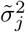 is the empirical variance of the log-transformed simulated measurements.

## Results and discussion

To demonstrate the properties of filter inference, we first infer posterior distributions from snapshot measurements for two modelling problems: 1. early cancer growth; and 2. EGF pathway signalling. We then compare the computational costs of NLME inference and filter inference and quantify the impact of using NUTS on the sampling efficiency. We conclude the section by highlighting the similarities between filter inference and summary statistics-based ABC and illustrate how inappropriate choices of filters may result in information loss or bias. Python scripts to reproduce the results are hosted on https://github.com/DavAug/filter-inference. All models are implemented in the open-source Python package `chi` [19], which we have extended to provide a user-friendly API for filter inference. For the inference, we use `pints`’ implementations of NUTS and the MH algorithm [29]. The gradients of the log-posterior, needed for NUTS, are automatically computed by `chi`, using the open-source Python package `myokit` [30].

### Early cancer growth model inference

To establish that filter inference is a sound approach for the inference of NLME models, we compare the results of filter inference and NLME inference on a common dataset. We synthesise snapshot measurements from the early cancer growth model, Eq 1, with a Gaussian error model, 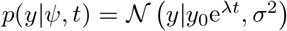, by first sampling individual-level parameters from the population model, 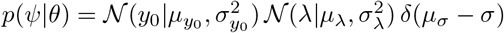. We then measure each individual by sampling from *p*(*y*|*ψ, t*). The population parameters are set to 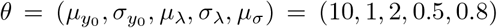 for the data-generation. 15 snapshot measurements are synthesised for 6 time points between 0 and 0.6 time units. The resulting dataset with a total of 90 measured individuals is illustrated by scatter points in Fig 3A. This dataset is still tractable for NLME inference. The details of the inference procedure and the convergence assessment are reported in S2 Appendix and S1 Table.

**Fig 3.**
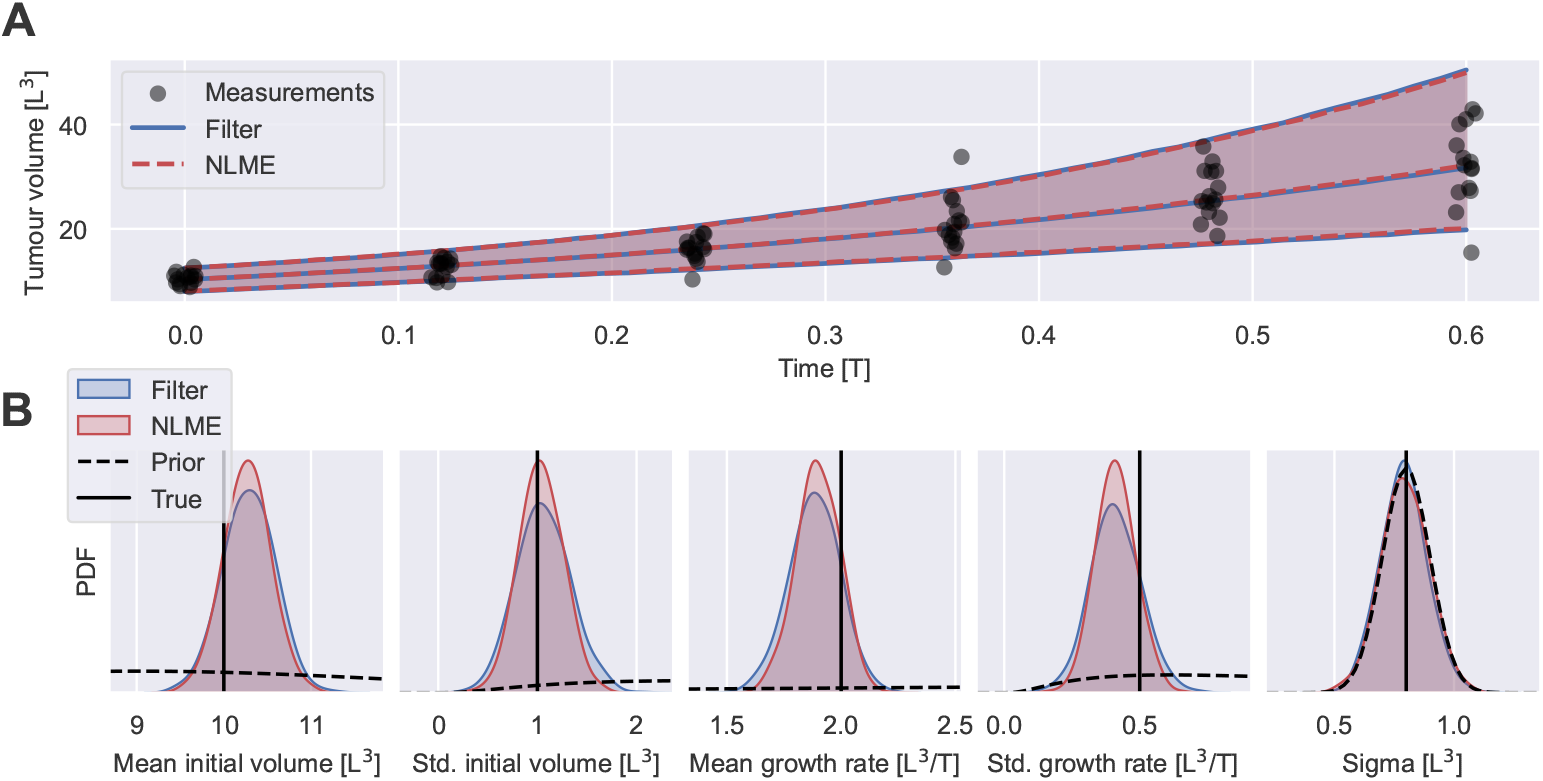
Filter inference versus NLME inference. A: Shows 90 snapshot measurements in arbitrary units, generated from the early cancer growth model, and the fitted NLME models obtained using filter inference with a Gaussian filter (blue) and NLME inference (red). The measurements are illustrated with jitter on the time axis. The fitted models are illustrated by the medians and the 5th to 95th percentile range of the inferred measurement distributions, 𝔼 _*θ*| 𝒟_ [*p*(*y θ, t*)]. The filter is constructed using *S* = 100 simulated individuals. B: Shows the inferred posterior distributions obtained using filter inference (blue) and NLME inference (red). The data-generating parameters (solid lines) as well as the prior distributions used for the inference (dashed lines) are also shown.

The inference results show that filter inference with a Gaussian filter and *S* = 100 simulated individuals and NLME inference produce almost identical fits to the measurements (see Fig 3A) and similar posterior distributions (see Fig 3B). Notably, the posterior distributions of both approaches encompass the data-generating parameter values within their main bulk probability mass. The measurements appear highly informative about the population means of the parameters but less so about the variability in the population. Importantly, the measurements are not informative about the noise parameter, *μ*_*σ*_, since the posterior distributions do not differ substantially from the prior distribution. This is because observed variability is not easily attributed to either IIV or measurement noise when individuals are not measured repeatedly. While the increase of the tumour volume variability over time indicates that IIV is present at least in one of (*y*_0_, *λ*), since a Gaussian error model cannot capture heteroscedasticity, the measurements in Fig 3A leave room for attributing all observed variability at *t* = 0 to just noise, or just IIV in *y*_0_. Thus, as long as their combined variability is of the same magnitude as the observed variability, each contribution may assume any magnitude between zero and the observed variability at *t* = 0. The variance of the measurements at *t* = 0 is 1.0 ± 0.4 (see S3 Appendix for the details of the estimation), while the prior of *μ*_*σ*_ focusses on values between 0.5 and 1 as shown in the right-most figure in Fig 3, constraining the noise variance to be at most of order 𝒪 (1). As a result, all values from the prior of *μ*_*σ*_ are compatible with the observations, and the posterior is not informed by the measurements. This lack of IIV-noise identifiability can be overcome when variability contributions from IIV and noise lead to distinct shapes of the measurement distribution, *p*(*y*|*θ, t*), and sufficiently many measurements are available to resolve such distributional differences. However, NLME inference from large snapshot datasets is computationally intractable and filter inference requires a large number of simulated individuals in order to distinguish IIV and noise, diminishing its computational advantages (see S4 Appendix for a detailed discussion). The efficient inference from snapshot measurements using filter inference with only a small number of simulated individuals is therefore reliant on informative noise priors. In many applications, such priors may be informed by the specifications of measurement devices. Where possible, repeated measurements of a few individuals may also be used to estimate noise parameters.

The inferred population distributions, represented by the averages over the posterior distributions, 𝔼 _*θ*| 𝒟_ [*p*(*ψ*|*θ*)], are illustrated together with the data-generating distribution in the left panel of Fig 4A (see S5 Appendix for details on the computation of 𝔼 _*θ*| 𝒟_ [*p*(*ψ*|*θ*)]). The comparison shows that the inference from 90 snapshot measurements with either inference method provides only a crude estimate of the IIV. This inaccuracy is a direct consequence of sampling bias – 90 individuals do not faithfully depict the whole population. To test whether this sampling bias can be mitigated by increasing the number of measured individuals, we exponentially increase the number of snapshot measurements per time point from 15 to 3 × 15 = 45, 3^2^ × 15 = 135 and 3^3^ × 15 = 405 snapshot measurements, resulting in datasets totalling *N* = 270, *N* = 810 and *N* = 2430 measurements. The inferred population distributions using filter inference are illustrated in panels 2, 3 and 4 of Fig 4A. The figure demonstrates that the estimation of the IIV improves with the number of measured individuals, as also quantified by the Kullback-Leibler (KL) divergence between the data-generating population distribution and the inferred distributions shown in Fig 4B. Overall, Fig 4 illustrates that many more than 90 snapshot measurements are needed to obtain accurate estimates of the IIV for the early cancer growth model. For higher dimensional NLME models, even more individuals are likely required, causing issues for NLME inference. In contrast, the computational costs of filter inference are not dominated by the number of measured individuals, and thus inference remains tractable.

**Fig 4.**
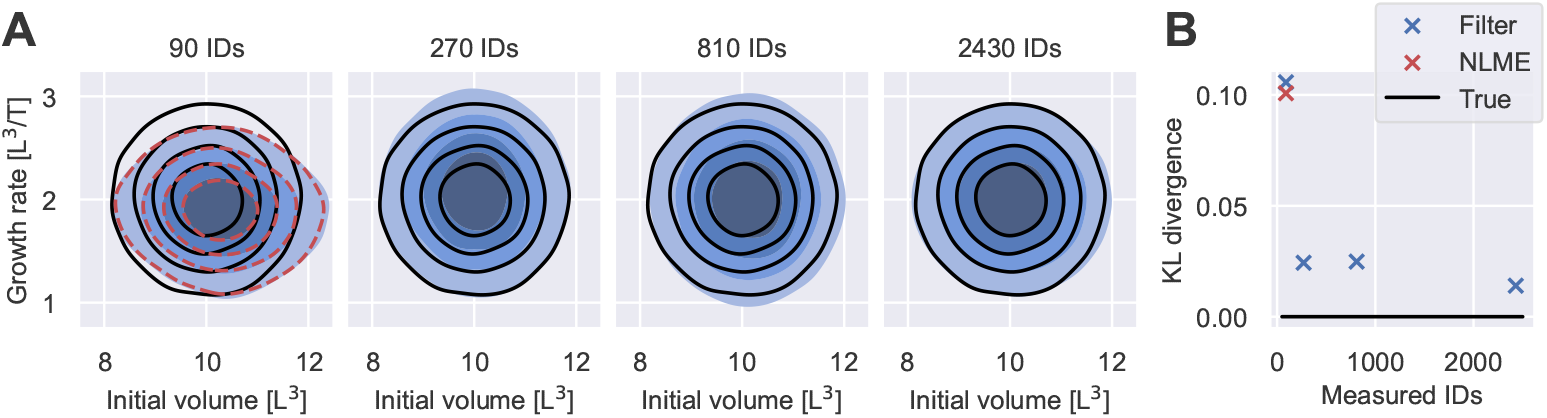
Quality of IIV estimates for varying dataset sizes. A: Shows inferred population distributions from measurements of varying numbers of individuals (IDs) using filter inference with a Gaussian filter and *S* = 100 simulated individuals in blue. The different shades of blue indicate the bulk 20%, 40%, 60% and 80% probability regions. The distribution inferred from measurements of 90 individuals using NLME inference from Fig 3 is illustrated by red dashed lines. The data-generating distribution is illustrated in black. B: Shows the KL divergences between the data-generating population distribution and the inferred distributions from A.

### EGF pathway model inference

The EGF pathway plays an important role in regulating the behaviour of epithelial cells and tumours of epithelial origin [31]. Understanding the cell-to-cell variability in the biochemical signalling is therefore of great interest [13]. Here, we demonstrate the ability of filter inference to estimate the parameters of a published EGF signalling pathway model from snapshot measurements.

In particular, we consider a model of inactive and active EGF receptor (EGFR) concentrations [32]

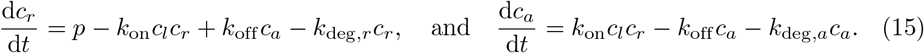

*c*_*r*_ models inactive and *c*_*a*_ active EGFR. The model assumes that inactive EGFR is produced at a constant rate, *p*. Upon binding of EGF, inactive EGFR is activated at a rate proportional to the surrounding EGF concentration *c*_*l*_. Once activated, receptors deactivate at a rate *k*_off_. Both, active and inactive receptors are assumed to degrade over time at rates *k*_deg,*a*_ and *k*_deg,*r*_, respectively. In our study, the cell-to-cell variability is modelled by varying production and activation rates. The remaining model parameters are fixed across cells. We synthesise two distinct datasets, each comprising 1200 cells and use the collective data from both datasets to perform filter inference. The first dataset contains snapshot measurements of cells exposed to a constant EGF concentration of *c*_*l*_ = 2 ng/mL, henceforth denoted as ‘data (low)’. The second dataset, ‘data (high)’, contains snapshot measurements of the same experiment with a higher constant EGF concentration of *c*_*l*_ = 10 ng/mL. The data are simulated using a lognormal error model centred on the model outputs with scale parameter *μ*_*σ*_. The measurements are generated over a period of 25 min with population parameters 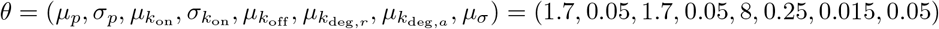, initialising both receptor concentrations at 0 ng/mL. The generated datasets are illustrated by black scatter points (data (low)) and grey scatter points (data (high)) in Fig 5A. We infer the model parameters from the synthetic datasets using Gaussian filters with *S* = 100 simulated cells. The noise parameter, *μ*_*σ*_, is fixed to the data-generating value during the inference. Details on the inference procedure are reported in S6 Appendix and S2 Table.

**Fig 5.**
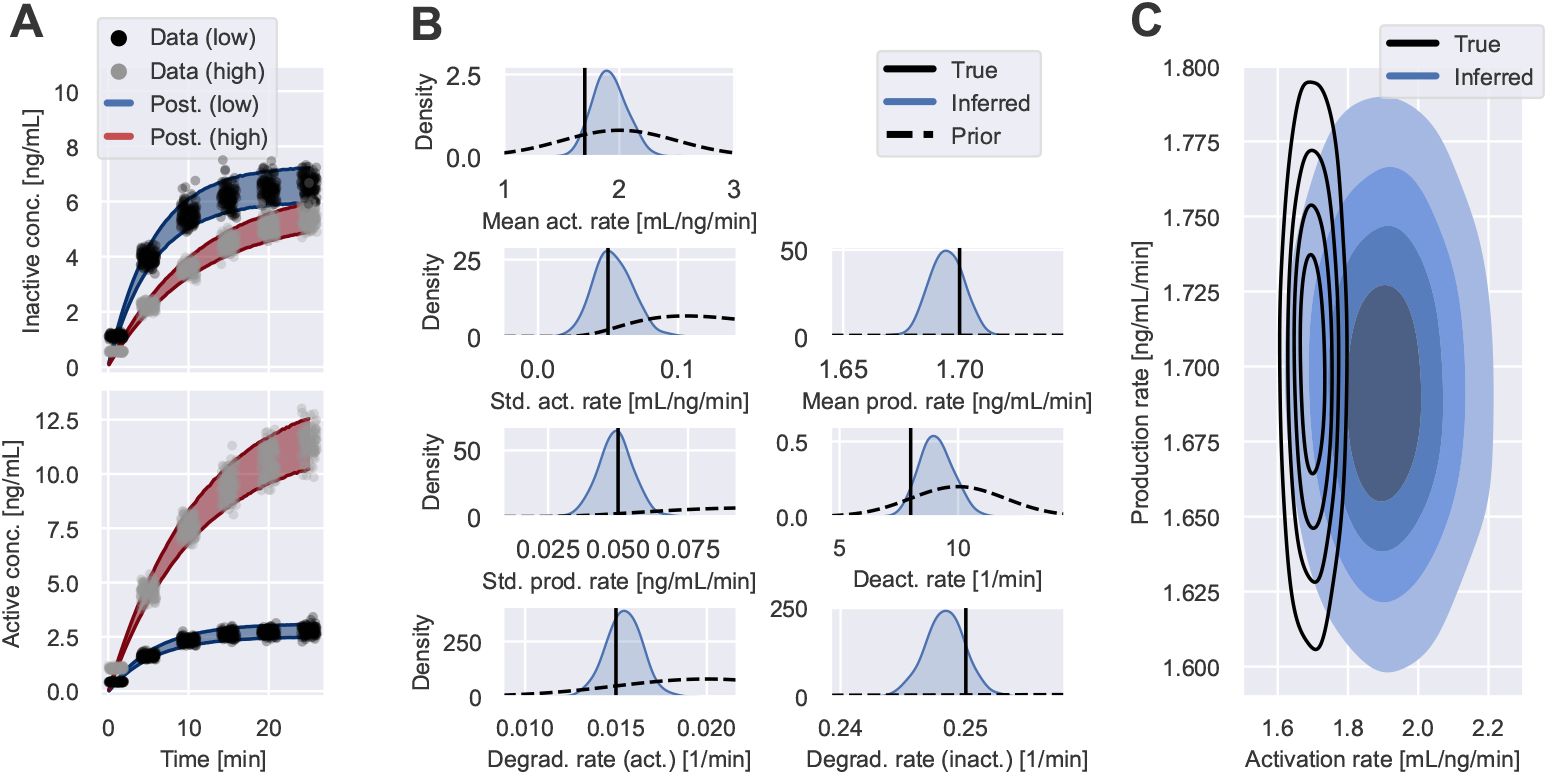
Inference results of EGF pathway model I. A: Shows snapshot measurements of active and inactive EGFR concentrations across cells. The cells are exposed to one of two EGF concentrations: data (low) with *c*_*l*_ = 2 ng/mL (black scatter points); and data (high) with *c*_*l*_ = 10 ng/mL (grey scatter points). The shaded areas illustrate the 5th to 95th percentile of the inferred measurement distributions using filter inference with Gaussian filters and *S* = 100 simulated cells. B: Shows the inferred posterior distributions of the model parameters illustrated by blue density plots, together with the data-generating parameter values illustrated by black solid lines. The density of the prior distribution is illustrated by dashed lines for each parameter. C: Shows the inferred population distribution of the production rate and the activation rate in blue. The different shades of blue indicate the bulk 20%, 40%, 60% and 80% probability regions. The probability regions of the data-generating distribution are illustrated by black contours.

Fig 5A shows that filter inference is able to infer measurement distributions that capture the dynamics of the observed EGF signalling pathway. Fig 5B shows that the inferred posteriors assign substantial weights to the data-generating parameters. The inferred cell-to-cell variability of the model parameters is of a reasonable magnitude, as the comparison of the data-generating population distribution and the inferred distribution in Fig 5C shows. However, the data appears to be more informative about the production rate variability than the activation rate variability. To investigate this further, in Fig 6A, we plot the posterior distribution of the population-level mean activation rate versus the deactivation rate. This indicates that it is not possible to identify both the activation rate and deactivation rates from the synthesised datasets. In Fig 6B, we show inference results when we fix the deactivation rate to its known fixed value – in this case, the inferred IIV distribution is closer to the true one.

**Fig 6.**
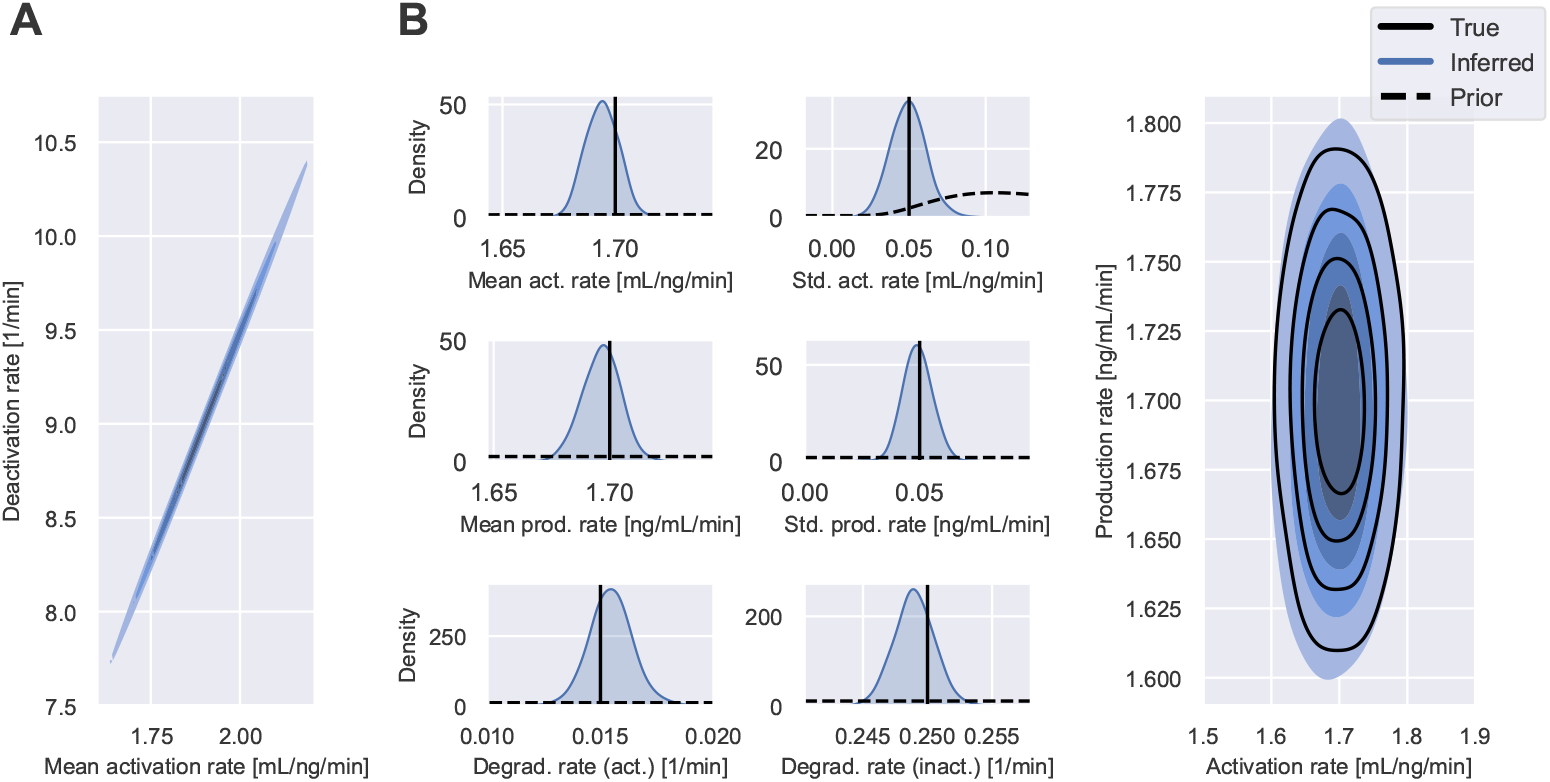
Inference results of EGF pathway model II. A: Shows the joint posterior distribution of the mean activation rate, 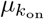, and the deactivation rate, 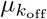 from Fig 5B. B: Shows the inferred posterior distribution and the inferred population distribution from a separate inference run, where we fixed the deactivation rate to its data-generating value. All other inference settings, including data and priors, remain unchanged from the inference approach used to generate Fig 5.

### Scaling & computational costs

Filter inference is able to infer the parameters of NLME models from thousands of snapshot measurements. This is because the dominant computational costs of the log-posterior evaluation do not scale with the number of measured individuals, as demonstrated Fig 7. The figure shows the evaluation time of the NLME log-posterior and its gradient (blue lines), defined in Eq 7, and the filter log-posterior and its gradient (red lines), defined in Eq 9, for the early cancer growth model and the EGF pathway model with increasing numbers of measured individuals. The filter log-posterior is defined using a Gaussian filter with varying numbers of simulated individuals. The gradients of the posteriors are automatically computed using chi [19]. Details on the estimation of the evaluation times are presented in S7 Appendix.

**Fig 7.**
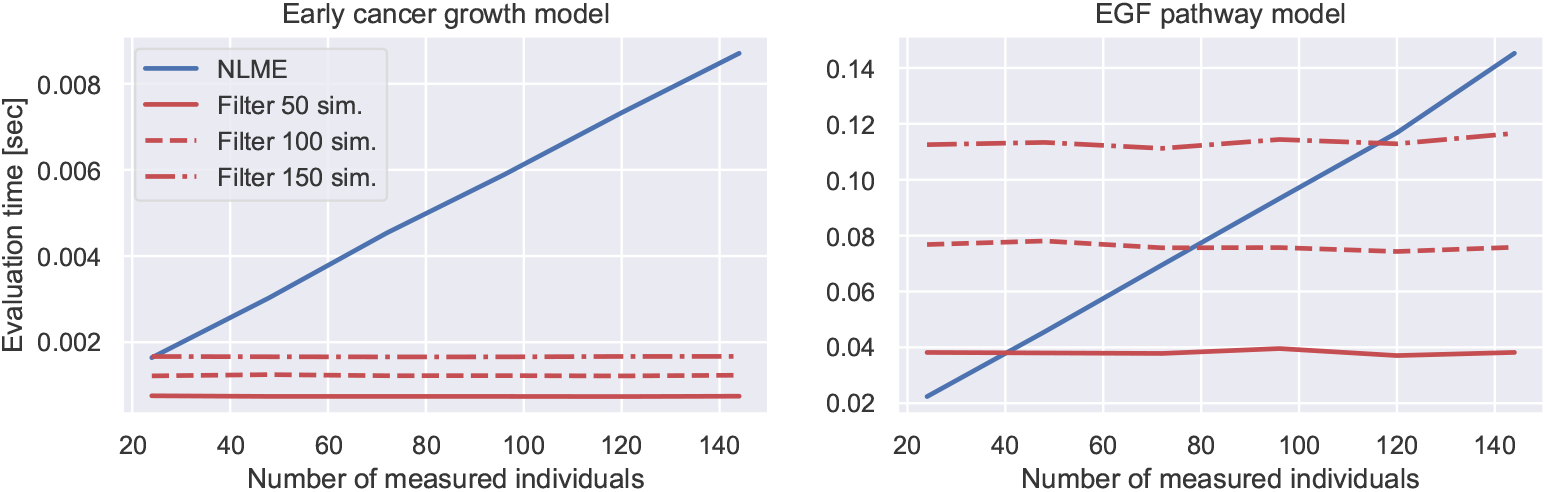
Computational costs of filter and NLME inference I - evaluation time of log-posterior. The figure shows the evaluation time of the NLME log-posterior and its gradient, defined in Eq 7, (blue lines) and the filter inference posterior and its gradient, defined in Eq 9, with a Gaussian filter and *S* = 50, *S* = 100 and *S* = 150 simulated individuals (red lines). The left and right panel illustrate the results for the early cancer growth model and the EGF pathway model, respectively.

The figure shows that the evaluation time of the NLME log-posterior scales linearly with the number of measured individuals, while the cost of the filter log-posterior remains constant. In particular, this illustrates that the computational costs of the filter log-posterior are roughly proportional to the number of simulated individuals, as discussed in the Methods. As a result, the speed up provided by filter inference is of order 𝒪 (*N/S*), where *N* and *S* denote the number of measured and simulated individuals, respectively. This reduces the log-posterior evaluation costs 34-fold for the early cancer growth model, and 13-fold for the EGF pathway model for datasets with 1000 snapshot measurements and filters with *S* = 100 simulated individuals. This cost reduction increases as the number of measured individuals grows.

However, in addition to the evaluation time of the log-posterior, the computational costs of inference are also determined by the total number of evaluations needed for convergence of the MCMC sampler. Since MCMC is a form of dependent sampling, there are typically autocorrelations between samples, reducing the number of i.i.d. samples drawn from the posterior distribution [33]. In order to determine the total computational costs of the approaches, we thus need to compare the number of log-posterior evaluations of NLME inference and filter inference needed for convergence.

Several metrics for the convergence assessment of MCMC samples exist, such as the 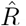-metric or the *R*^*^-metric [34, 35]. These metrics could be used to determine the number of log-posterior evaluations needed to reach a certain degree of convergence for different MCMC algorithms. An alternative approach to compare the computational costs of NLME inference and filter inference is to infer the posteriors using an MCMC algorithm that uses an initial calibration phase to adjust the number of log-posterior evaluations per MCMC step in order to maximise the convergence rate across inference problems. Such an algorithm is NUTS [17, 18]. Using its calibration strategy, NUTS converges within 1000 MCMC iterations post calibration for the early cancer growth model and the EGF pathway model. We choose to estimate the total computational costs of NLME inference and filter inference using the latter approach, and run 1500 NUTS iterations for both, the early cancer growth model and the EGF pathway model for different datasets with varying numbers of measured individuals (see Fig 8). The initial 500 iterations of each inference run are used to calibrate the algorithm. Each evaluation includes the evaluation of the log-posterior and its gradient.

**Fig 8.**
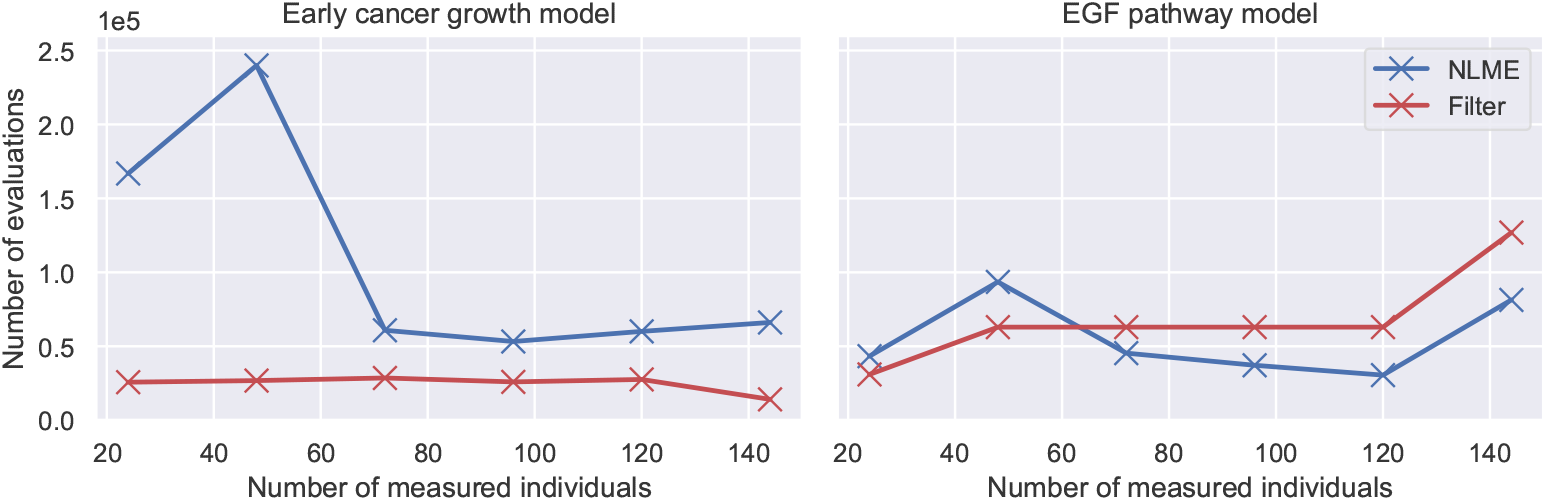
Computational costs of filter and NLME inference II - number of log-posterior evaluations. The figure shows the number of log-posterior evaluations of NLME inference (blue lines) and filter inference, with a Gaussian filter and *S* = 100 simulated individuals, (red lines) for the early cancer growth model and the EGF pathway model using varying sizes of snapshot datasets. Each log-posterior evaluation includes the evaluation of its gradient. The posterior distributions are inferred using 1000 MCMC iterations of NUTS after calibrating the algorithm for 500 iterations.

The figure shows that the number of log-posterior evaluations of NLME inference and filter inference are of the same order of magnitude for the investigated models. Each log-posterior evaluation includes the evaluation of the log-posterior gradient. For the early cancer growth model, NUTS requires fewer evaluations during filter inference, while for the EGF pathway model NUTS evaluates the log-posterior less often during NLME inference. Overall, our analysis demonstrates that the benefit to using filter inference scales linearly with the number of measured individuals, meaning, for large datasets typical in cell biology, it can be orders of magnitude faster than NLME inference.

### Sampling efficiency

Filter inference can reduce the costs of inference in both its stochastic and its differentiable form. In this section, we investigate the degree to which the differentiable version and the use of NUTS improves the efficiency of filter inference.

To this end, we estimate the sampling efficiency of the approaches using the effective sample size (ESS) metric, defined in [36]. The ESS estimates the number of i.i.d. samples drawn from each posterior dimension by an MCMC algorithm. Thus, the larger the ESS for a fixed number of log-posterior evaluations, the better the sampling efficiency of the algorithm. In Fig 9 we show the minimum ESS across dimensions, obtained from inferring the early cancer growth model posterior and the EGF pathway model posterior. We infer the posteriors twice: 1. using the MH algorithm and the stochastic version of filter inference, defined in Alg 2; and 2. using NUTS and the differentiable version of filter inference, defined in Alg 3.

**Fig 9.**
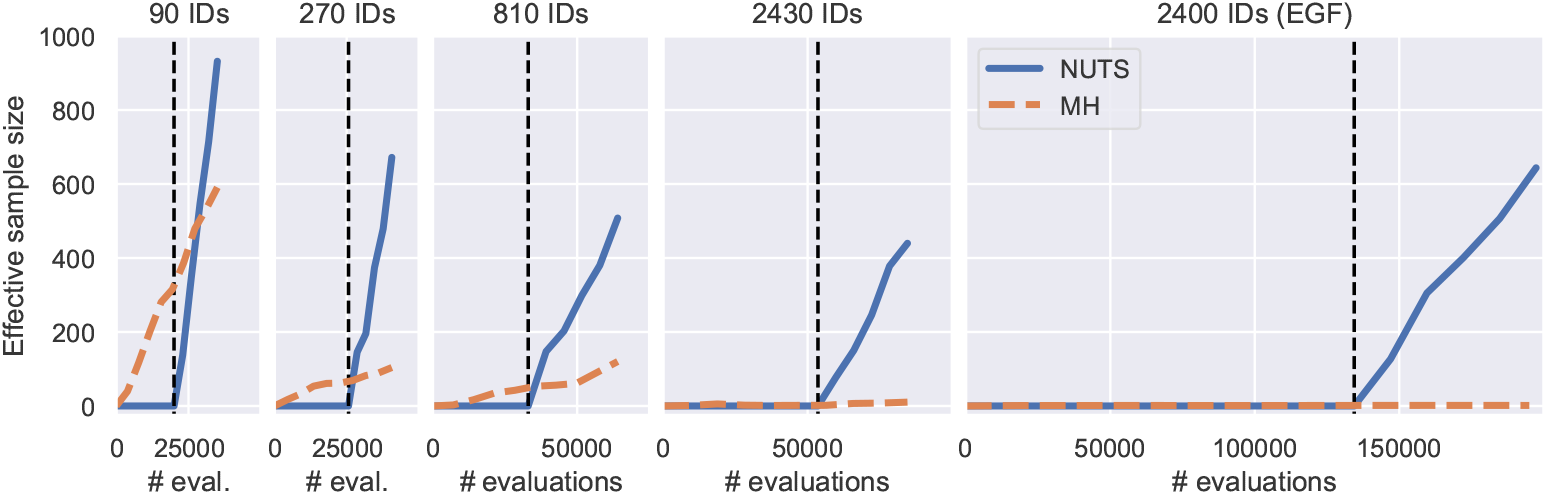
Sampling efficiency of filter inference variants. The figure shows the minimum ESS across dimensions as a function of log-posterior evaluations for different posterior distributions inferred with filter inference using 2 MCMC algorithms: 1. NUTS (blue); and 2. MH (orange). For NUTS, the number of evaluations include evaluations of the log-posterior gradient. For MH, the log-posterior gradient is not evaluated. Panels 1, 2, 3 and 4 show the minimum ESS of the cancer growth model posteriors from Fig 4 and panel 5 shows the EGF pathway model posteriors from Fig 6B.

The figure shows that the sampling efficiency is improved when using NUTS. After 35,000 iterations, the MH algorithm generated an ESS of approximately 600 when the posterior was inferred from the cancer growth dataset with 90 measured individuals (see left panel in Fig 9), while NUTS generated an ESS of 932 after 1500 iterations, using the first 500 iterations for calibration. These 1500 iterations translate into 34,985 log-posterior and gradient evaluations (see S3 Table) out of which the majority (20,000 evaluations) are from the calibration phase and therefore do not contribute to the ESS. Evaluating the gradient in addition to the log-posterior approximately doubles the evaluation costs (see S4 Fig). As a result, NUTS generated an ESS that is 1.5 times greater with approximately the same (2 × 14, 985*/*35, 000 ≈ 1) computational costs post calibration. This efficiency advantage becomes more pronounced as the number of measured individuals increases. In particular, the ESS of the MH algorithm, despite extensive efforts to tune its hyperparameters (see S8 Appendix for details), is smaller than 10 after 90,000 MH iterations for both the early cancer growth model and the EGF pathway model, when datasets with thousands of measurements are used for the inference (see panels 4 and 5 in Fig 9). At the same time, NUTS is able to generate an ESS of order 100 for all datasets and models by automatically tuning the number of evaluations per MCMC step during the calibration phase. The number of log-posterior evaluations across all inference runs are reported in S3 Table.

Our study confirms that NUTS and Alg 3 perform better across inference problems than the MH algorithm and Alg 2. In addition, our study indicates that it is more challenging to sample from filter posteriors when more individuals are measured (see ESS per evaluation in Fig 9). NUTS is nevertheless able to achieve good sampling efficiencies by adaptively using more log-posterior evaluations during the calibration phase. With the stochastic version of filter inference, on the other hand, we were not able to achieve comparable sampling rates when many snapshot measurements were measured. To understand this behaviour, we investigate the relationship between filter inference and summary statistics-based ABC in the next section.

### Relationship to summary statistics-based ABC

Filter inference conceptually extends the ABC inference framework, as discussed in the Methods. While ABC is a hugely successful inference strategy across fields, such as population genetics [37], epidemiology [38] and climate modelling [39], a common criticism of ABC is the need to manually choose error margins in order to quantify the distance between summary statistics of real and simulated measurements. Too large error margins reduce the quality of the inference results, while too narrow error margins lead to high rejection rates and, thus, to a poor sampling efficiency. A theoretically justified choice for error margins reflects the finite sampling variation of the data summary statistics relative to the exact summary statistics of the data-generating distribution. In contrast, filter inference does not define error margins, despite some filters being defined by summary statistics. A Gaussian filter, for example, is defined by the mean and variance of the simulated measurements. Denoting such filters henceforth as summary statistics-based (SS-based) filters, we may expect filter inference with SS-based filters to behave similarly to ABC based on the same summary statistics. In Fig 10 we investigate this intuition by revisiting the filter inference results from Fig 4.

**Fig 10.**
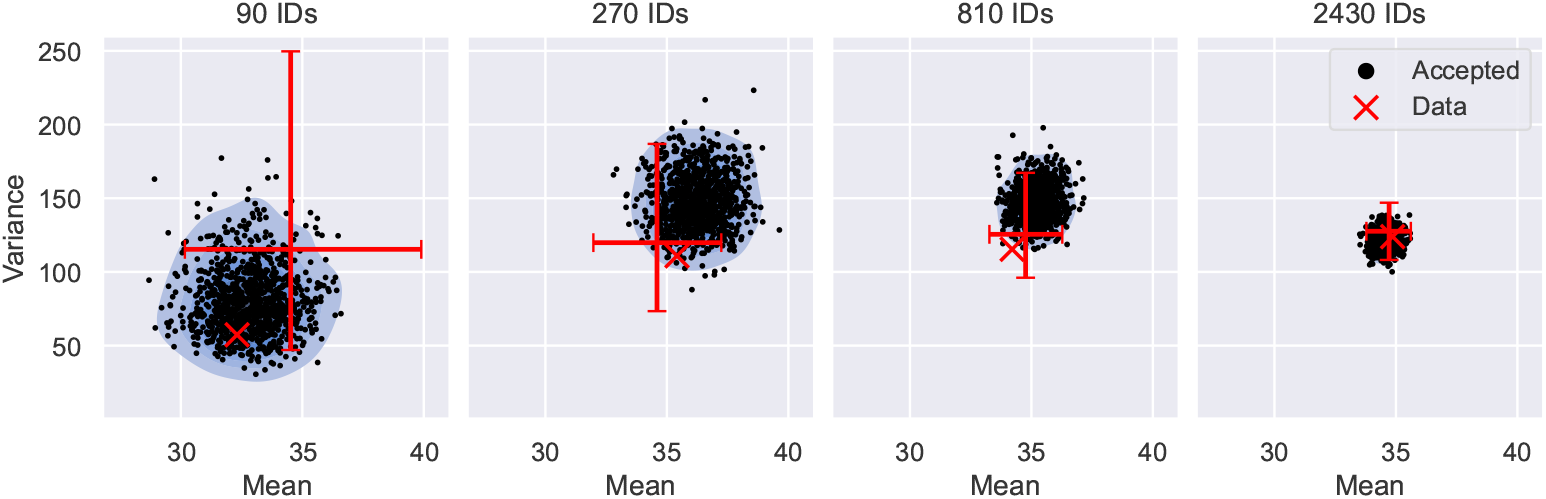
ABC interpretation of filter inference. The figure shows the accepted means and variances of the Gaussian filter at *t* = 0.6 from the inference results in Fig 4, where filter inference is performed on datasets with 90, 270, 810 and 2430 snapshot measurements of the cancer growth model. The accepted summary statistics are illustrated as black scatter points with the corresponding KDE plots shown in blue. The summary statistics of each dataset is illustrated by a red scatter point. The sample variation of the dataset summary statistics is represented by the 5th to 95th percentile of the summary statistic distribution (red bars), estimated from 1000 realisations of each dataset. The exact mean and variance of the data-generating distribution is close to the intersection of the bars.

The figure shows the accepted means and variances at time *t* = 0.6 (black points) of the cancer growth inference for datasets of different sizes. As a reference, the means and variances of the datasets used for the inference are illustrated as red crosses in the figure. To illustrate the sample variation of the data summary statistics, we generate 1000 realisations of each dataset from the data-generating distribution and plot the 5th to 95th percentile of the resulting summary statistic distribution as red bars. The exact mean and variance of the data-generating distribution are approximately equal to the value marked by the intersection of the red bars. The figure shows that filter inference accepts simulated summary statistics within an error margin around the data summary statistics, notably without explicitly computing the data summary statistics during the inference. The magnitude of the error margins scales with the number of measured individuals and is comparable to the 5th to 95th percentile interval of the data summary statistics across dataset sizes.

These observations suggest that filter inference with a Gaussian filter is similar to a special variant of ABC based on the mean and variance, where the error margin is automatically scaled with the uncertainty of the summary statistic estimates. This similarity is a consequence of the Gaussian filter construction, defined in Eq 10. The Gaussian filter likelihood assigns high likelihood only to simulated means and variances that are compatible with the data. When the dataset contains few observations, the filter likelihood is flat and permits large deviations from the mean and variance of the data, while datasets containing more observations become more restrictive. This leads to an automatic scaling of the error margin with the uncertainty of the data summary statistics. In S9 Appendix, we provide a proof that in the limit *N* → ∞ filter inference with a Gaussian filter is equivalent to ABC based on the mean and variance with vanishing error margins. In S10 Appendix, we further prove that this equivalence also extends to other SS-based filters, provided the filter likelihood has a unique and identifiable maximum and the maximum likelihood estimates (MLEs) converge to the summary statistics of the data.

Revisiting the results in Fig 9, Fig 10 also provides an explanation of why the number of measured individuals reduces the ESS for the stochastic approach. Alg 2 simulates measurements, and thus summary statistics, for each proposal randomly. The estimation error of these simulated summary statistics is determined by the number of simulated individuals. The error margin around the data summary statistics, on the other hand, is set by the number of measured individuals. As a result, proposals close or equal to the data-generating parameters may nevertheless be randomly rejected when the error margin is smaller than the estimation error of the simulated summary statistics. For a fixed number of simulated individuals, this rejection rate becomes larger as the number of measured individuals increases, resulting in a poor sampling efficiency (see Fig 9). In contrast, the gradient-based sampling strategy of NUTS, together with Alg 3, avoids rejections due to the randomness of simulated measurements entirely, providing high sampling efficiencies also for large numbers of measured individuals.

### Information loss and bias

In contrast to NLME inference, filter inference is an approximate inference approach. This approximation may result in information loss and bias. The potential for inaccurate inference results is common to all ABC methods and, in this case, comes from the filter approximation of the population-level log-likelihood, Eq 8. Filters construct a noisy estimate of the likelihood from measurements of a small number of simulated individuals. The fewer individuals are simulated, the lower the costs of the log-likelihood evaluation. This provides an incentive to reduce the number of simulated individuals as much as possible. However, as illustrated in Fig 11 for the early cancer growth model, there is a trade-off between computational costs and the accuracy of the IIV estimation. The fewer individuals are simulated, the more information is lost about the variability in the population. The optimal number of simulated individuals will likely vary between problems and modelling rationales. However, in our experiments we achieve reasonable results with *S* = 100 simulated individuals across models.

**Fig 11.**
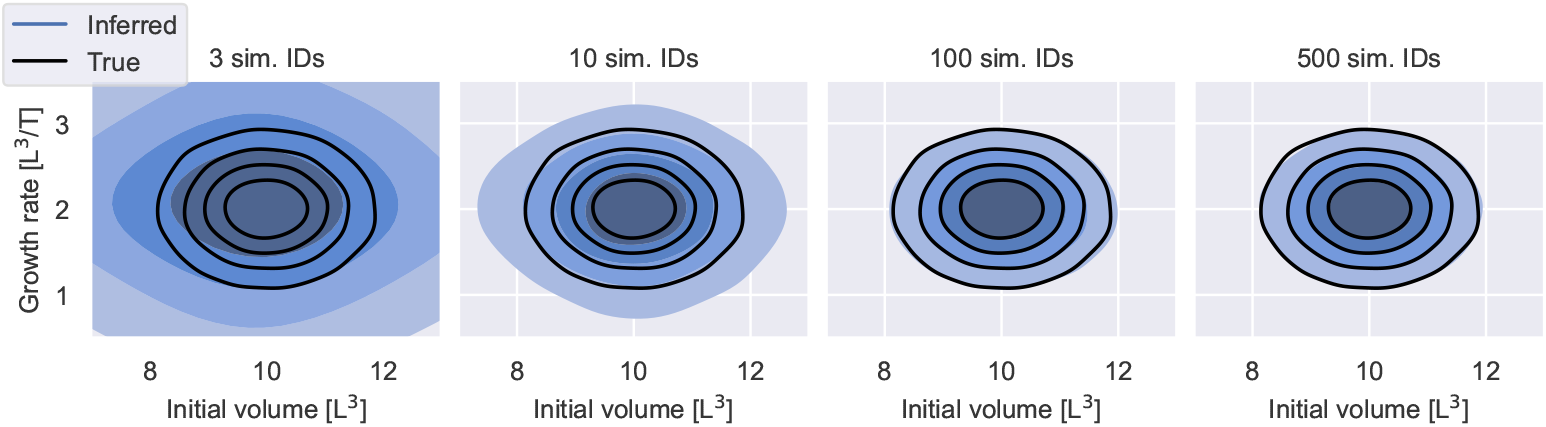
Information loss I - number of simulated individuals. The figure shows inferred population distributions from 2430 snapshot measurements of the early cancer growth model, using filter inference with a Gaussian filter and *S* = 3, *S* = 10, *S* = 100 and *S* = 500 simulated individuals. The inferred distributions are visualised in blue. The different shades of blue indicate the bulk 20%, 40%, 60% and 80% probability regions. The data-generating distribution is illustrated by black contours.

A second source for information loss and bias is the choice of the filter itself. To illustrate this effect we modify the cancer growth model to two patient subpopulations with different variants of cancer: an aggressive variant and a moderate one. This is implemented by defining a covariate-dependent mean growth rate, *μ*_*λ*_(*x*) = *μ*_*λ,m*_ +*χ* Δ*μ*_*λ*_, where *χ* = 0 indicates patients with the moderate variant and *χ* = 1 patients with the aggressive variant, resulting in a multi-modal population distribution

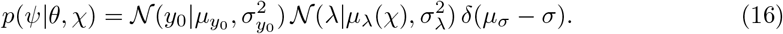

We synthesise two datasets: one dataset with snapshot measurements from 120 individuals; and one with measurements from 3000 individuals. In both cases, half of the individuals have the aggressive cancer variant and the other half the moderate variant. The data are synthesised with 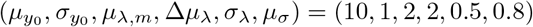.

The inference results for different choices of filters and *S* = 100 simulated individuals are illustrated in Fig 12. Where tractable, NLME inference is used to infer the exact posterior distribution. Otherwise, the data-generating distribution is used as a reference for the inference results. The figure shows that the quality of the results varies substantially with the choice of the filter. The Gaussian and lognormal filters yield reasonable approximations of the overall individual-level variability, but are not able to resolve the multi-modal structure of the growth rate when only 120 patients are measured. For 3000 measured patients, both filters begin to distinguish the moderate and aggressive cancer growth subpopulations. In comparison, inference with a Gaussian mixture filter with two kernels resolves the multi-modal population structure for both numbers of measured individuals (see middle panel in 12). Inference with a Gaussian KDE filter or a lognormal KDE filter similarly resolves the multi-modal population structure, but, here, the filter posteriors underestimate the IIV for both numbers of measured individuals.

**Fig 12.**
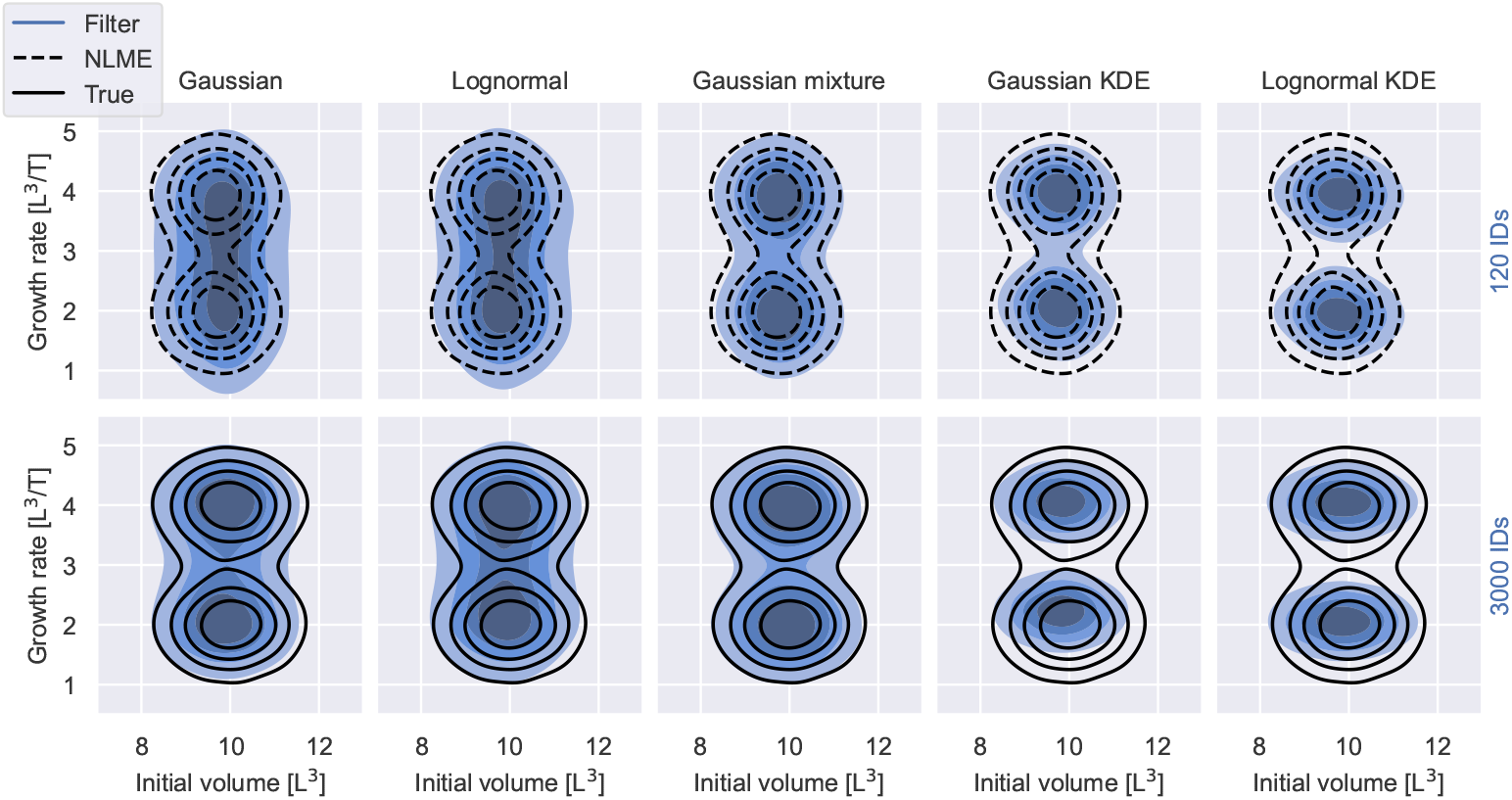
Information loss II - choice of filter. The figure shows inferred population distributions from 120 (top row) and 3000 (bottom row) snapshot measurements, using filter inference with different filter choices and *S* = 100 simulated individuals. The inferred distributions, 𝔼 _*θ*| 𝒟_ [*p*(*ψ θ*)], are visualised in blue. The different shades of blue indicate the bulk 20%, 40%, 60% and 80% probability regions. The data-generating distribution is illustrated by black solid lines. The posterior inferred with NLME inference is illustrated by black dashed lines.

The reasons for the observed information loss and biases are different for SS-based filters and KDE-based filters. Intuitively, it is not surprising that SS-based filters with a single mode, such as the Gaussian filter and the lognormal filter, may produce inaccurate inference results when the true measurement distribution is multi-modal. We nevertheless observe in Fig 12 that both filters resolve the multi-modal structure in the population distribution when 3000 individuals are measured. To understand this behaviour, recall that filter inference with SS-based filters is similar to ABC (see Relationship to summary statistics-based ABC) and that ABC converges to the true posterior distribution in the limit where 1. the error margin of the summary statistics goes to zero; 2. the summary statistics are sufficient; and 3. the number of simulated measurements is the same as the number of observed measurements [40, 41]. Filter inference, on the other hand, can be equivalent to ABC for certain SS-based filters in the limit where *N* → ∞ and the error margin of ABC vanishes (see S10 Appendix). Filter inference may therefore converge in the limit where 1. the number of measured individuals goes to infinity; 2. the filters are defined by sufficient summary statistics; and 3. the number of simulated measurements is the same as the number of observed measurements. This may explain why the Gaussian and the lognormal filters start to resolve the multi-modal population structure for 3000 measured individuals, but not for 120 measured individuals. To support this intuition, we compare the filter inference results from Fig 12 to the true measurement distribution in Fig 13. The left panel shows the histogram over all accepted simulated measurements at *t* = 0.6 during the inference with the Gaussian filter (blue) for the dataset with 3000 measured individuals and the corresponding data-generating measurement distribution (black). The histogram over simulated measurements captures the multi-modal structure of the data-generating measurement distribution, despite the use of a Gaussian filter. A typical filter, sampled during the inference, is illustrated in red.

**Fig 13.**
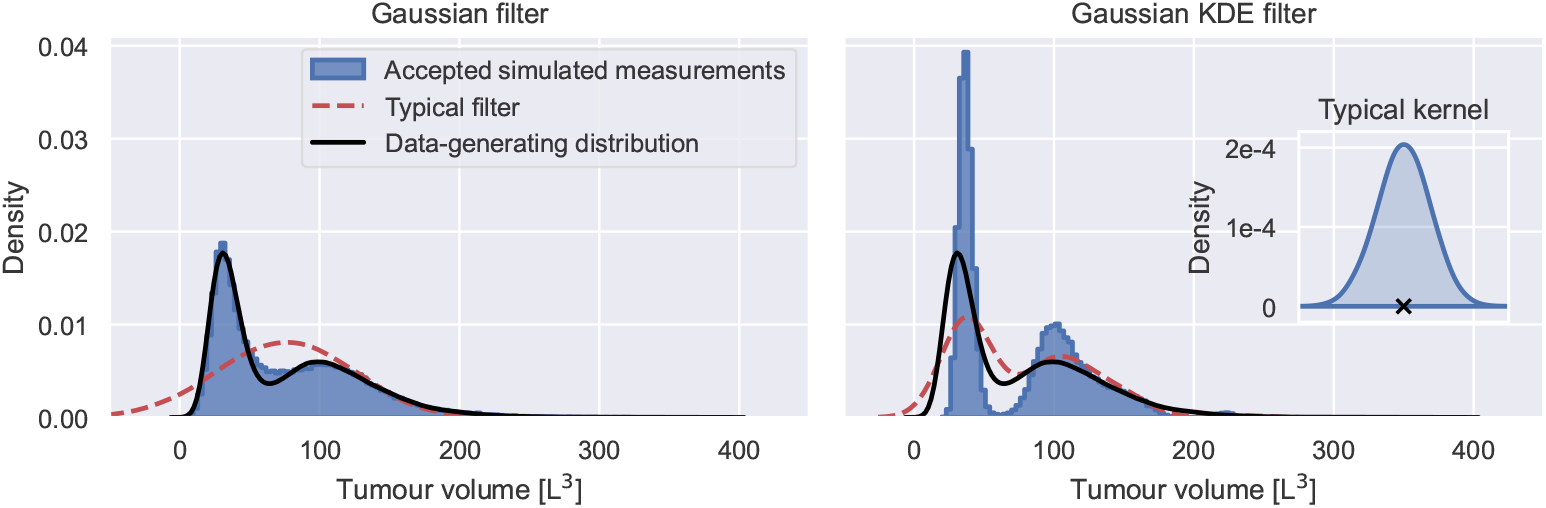
Accepted simulated measurements during filter inference. The figure shows the histograms over all accepted simulated measurements at *t* = 0.6 (blue) during inference with a Gaussian filter (left panel) and a Gaussian KDE filter (right panel) from the bottom panel in Fig 12. The data-generating measurement distribution is illustrated in black. A typical realisation of each filter is illustrated in red. The inset figure in the right panel shows a typical kernel (blue) placed on a simulated measurement (black cross) during the construction of the Gaussian KDE filter. The simulated measurement is placed at a tumour volume of 350 for illustration purposes. The scale of the kernel is taken from the filter realisation.

KDE-based filters, on the other hand, cannot be interpreted in the same way. To understand the behaviour of KDE-based filters, note that the filter log-likelihood is linked to the KL divergence between the filter and the observed measurement distribution

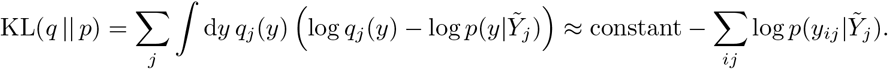

Here, *q*_*j*_ denotes the data-generating distribution at time *t*_*j*_, which we approximate by the observed measurements on the right hand side. We identify the last term on the right as the negative log-likelihood of the filter (see Eq 8). Consequently, the filter likelihood is maximised when the filter minimises the KL divergence, i.e. the filter optimally approximates the observed measurement distribution. For KDE-based filters, this leads to a bias of accepting simulated measurements that are closer to high density regions of the observed measurement distribution during inference, as illustrated in the right panel of Fig 13. The figure shows that the histogram over all accepted simulated measurements (blue) during the inference with the Gaussian KDE filter with *S* = 100 simulated individuals does not faithfully represent the data-generating distribution (black), despite typical filters (red) providing reasonable approximations. This is because KDE-based filters approximate the measurement distribution by averaging the densities of *S* finite-sized kernels placed on each of the simulated measurements (see Eqs 13 and 14). The size of the kernels extends the reach of simulated measurements, here approximately 50 tumour volume units in both directions (see inset figure). This makes it unnecessary and even unfavourable to occupy low density regions of the observed measurement distribution, resulting in a bias towards high density regions. The underrepresentation of simulated measurements in low density regions of the observed measurement distribution leads to an underestimation of the IIV, as demonstrated in Fig 12.

However, as the number of simulated measurements tends to infinity, the size of the kernels goes to zero (see Eqs 13 and 14) and the bias towards high density regions vanishes. If the observed measurement distribution is identical to the data-generating distribution, this implies convergence to the data-generating parameters. But, if the observed measurement distribution is not representative for the whole population, KDE-based filters will overfit the observed distribution, and thus filter inference will underestimate the variability in the population and the uncertainty in the parameter estimates.

In practice, inference is performed on datasets with a finite number of measured individuals using a finite number of simulated individuals. In this context, SS-based filters appear to provide a better accuracy-cost trade-off, especially when informed summary statistic choices are possible, as the middle column of Fig 12 demonstrates. Here, we infer the population distribution from the synthesised datasets using a Gaussian mixture filter with *S* = 100 simulated individuals and two kernels. Two Gaussian kernels are able to represent the bimodal structure of the observed measurement distribution more faithfully, resulting in inference results with negligible information loss.

## Conclusion

Filter inference is an efficient and scaleable inference approach for NLME models, enabling the study of variability from previously intractable datasets, for example, snapshot measurements of potentially thousands of individuals. However, filter inference also introduces new challenges, such as the potential for information loss and bias, which currently can only be understood with the help of repeated synthetic data-generation and inference cycles. In contrast to other snapshot inference methods, the efficiency and scalability of filter inference also centrally relies on the availability of gradients, which may be difficult to obtain for large systems of differential equations.

## Supporting information

**S1 Appendix. Optimised implementation of filter inference**. The implementation of filter inference can be optimised such that it solves the time series model only once for each simulated individual by leveraging the fact that numerical differential equation solvers can efficiently evaluate a system of differential at any number of time points. The optimised implementation, therefore, solves the time series model for each individual only once up front, computing 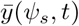 for all measured time points in one time series model solve. This removes the unnecessary costs of integrating the time series model for each measured time point separately.

**S2 Appendix. Estimation of early cancer growth model parameters**. The parameters of the early cancer growth model are estimated with pints’ implementation of NUTS, unless otherwise specified. MCMC chains are run for 1500 iterations, where the first 500 iterations are used for calibration. The convergence is assessed using the 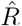 -statistic from 3 MCMC chains, initialised at randomly sampled points from the prior distribution. The prior distribution for all inference runs is

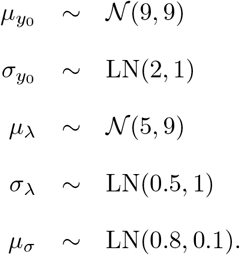

Detailed information on the convergence can be found in S1 Table.

**S3 Appendix. Estimation of variance estimation error**. We estimate the variance at time point *t*_*j*_ using

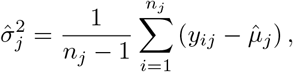

where *n*_*j*_ denotes the number of measurements at *t*_*j*_ and 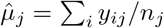 denotes their mean. Assuming that *y*_*ij*_ are drawn from a Gaussian distribution 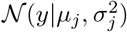 with mean *μ*_*j*_ and variance 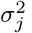, the normalised variance estimator follows a *χ*^2^-distribution with *n*_*j*_ − 1 degrees of freedom

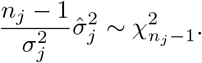

The variance of a 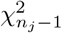 random variable is 2(*n*_*j*_ − 1). As a result, we can estimate the standard deviation of 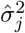 using

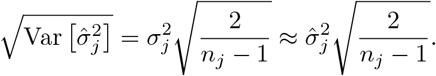

At *t* = 0, the measurement distribution in Fig 3A happens to be Gaussian, and the above estimate is unbiased.

**S4 Appendix. IIV-noise distinguishability**. To understand the IIV-noise distinguishability when estimating parameters from snapshot measurements using NLME inference and filter inference, we generate snapshot datasets from the early cancer growth model analogously to Early cancer growth model inference, but with increased numbers of measured individuals per time point. In particular, we generate datasets with *N* = 90, *N* = 270, *N* = 810 and *N* = 2430 individuals and estimate the parameters using NLME inference and filter inference with a Gaussian filter and *S* = 100 simulated individuals. The inference results are depicted in S1 Fig.

The figure shows that the uncertainty of the population mean estimates and the population standard deviation estimates decreases as the number of measured individuals increases, demonstrating that the estimation of IIV becomes more accurate as more individuals are measured. However, it also shows that for most of the datasets the posterior distributions of the noise parameter, *μ*_*σ*_, do not substantially differ from the prior distribution. As argued in Early cancer growth model inference, this is because the prior distribution, *p*(*μ*_*σ*_), focuses on values that give rise to noise magnitudes that are all compatible with the variability of the measurements, such that variability contributions from IIV and noise can only be distinguished if they give rise to different shapes of the measurement distribution, *p*(*y*|*θ, t*), and sufficiently many measurements are available to resolve such distributional differences. S2 Fig shows the measurement distribution for different samples from the posterior distribution in the top row of S1 Fig, demonstrating that the shape of the distribution does change for different contributions from IIV and noise, while the variance of the distribution remains the same.

Such distributional differences cannot be resolved by *N* = 90, *N* = 270 and *N* = 810 snapshot measurements, as demonstrated by columns 1-3 in S1 Fig. However, using snapshot measurements from *N* = 2430, NLME inference begins to have sufficient statistical power to resolve differences between IIV and noise, resulting in an update of *p*(*μ*_σ_| 𝒟) relative to the prior (see bottom right corner of S1 Fig). Filter inference with a Gaussian filter and *S* = 100 simulated individuals is not able to distinguish IIV and noise, even when *N* = 2430 individuals are measured. This is a consequence of the filter approximation of the measurement distribution which only uses simulations of *S* = 100 individuals. While *N* = 2430 measurements start to resolve distributional differences between IIV and noise contributions, *S* = 100 simulated individuals are insufficient to do so. Increasing the number of simulated individuals alleviates this limitation, as demonstrated in S3 Fig, where we illustrate posterior distributions inferred from *N* = 60 000 snapshot measurements using filter inference with Gaussian filters and *S* = 5 000 simulated individuals.

**S5 Appendix. Estimation of posterior average of population distribution**. Formally, the average of the population distribution over the posterior distribution, also known as the posterior predictive population distribution, is defined as

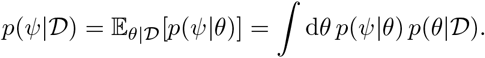

While the integral is generally difficult, or impossible, to compute analytically, the above equation implicitly defines a joint distribution of the individual-level parameters and the population-level parameters, *p*(*ψ, θ*|D) = *p*(*ψ*|*θ*) *p*(*θ*| 𝒟). To sample from *p*(*ψ*| 𝒟), we can sample from *p*(*ψ, θ*| 𝒟) and marginalise over the sampled population-level parameters afterwards. In particular, we first sample a *θ* from *p*(*θ*| 𝒟), and then draw a *ψ* from *p*(*ψ*|*θ*) using the sampled *θ*. The histogram over the sampled *ψ* converges to the posterior predictive population distribution as the number of samples tends to infinity. In practice, samples from *p*(*θ*| 𝒟) are generated by sampling with replacement from the MCMC posterior samples.

**S6 Appendix. Estimation of EGF pathway model parameters**. The parameters of the EGF pathway model are estimated with pints’ implementation of NUTS, unless otherwise specified. MCMC chains are run for 1500 iterations, where the first 500 iterations are used for calibration. The convergence is assessed using the 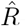 statistic from 3 MCMC chains, initialised at randomly sampled points from the prior distribution. The prior distribution for all inference runs is

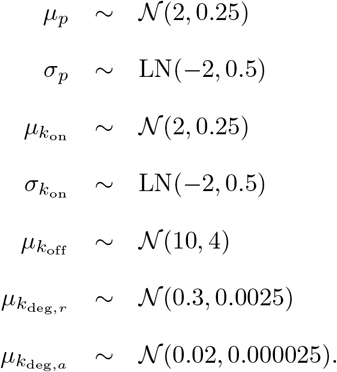

Detailed information on the convergence can be found in S2 Table.

**S7 Appendix. Evaluation time estimation**. The evaluation times of the NLME log-posterior and the filter log-posterior in Fig 7 are estimated by evaluating the log-posteriors for each dataset 10 times at fixed parameter values on a MacBook Pro, 2.8 GHz Quad-Core Intel Core i7 processor with 16 GB 2133 MHz LPDDR3 memory. The execution time of each evaluation is measured using the Python package timeit. The minimum execution time of the 10 repeats is reported in the figure. In this way, we are able to find the optimal execution time in Python, accounting for Python’s unpredictable overhead of execution time.

**S8 Appendix. Hyperparameter tuning of MH algorithm**. The hyperparameter of the MH algorithm is the step size of the proposal step. More precisely the MH algorithm proposes new parameters by sampling from a multivariate Gaussian distribution centred at the last accepted proposal. The covariance matrix of this distribution sets the scale of the step size and is the hyperparameter of the algorithm. To find a good set of step sizes we performed a grid search over several step sizes (see S4 Table). In particular, we used the variances of the filter posterior distributions inferred using NUTS in S1 Fig and Fig 6B as a starting point – information one would usually not have available prior to the inference. This allowed us to carry out the grid search more efficiently, as the marginal variances of the target distribution provide a good starting point for the diagonal of the covariance matrix of the MH proposal distribution. For each problem we ran the MH algorithm 7 times with 3 chains for 5000 MCMC iterations with diagonal covariance matrices whose diagonal was set to *x* times the variances of the target distribution. We performed the grid search for *x* ∈ {1.4, 1.2, 1, 0.8, 0.6, 0.4, 0.2}. The covariance matrix with the largest ESS was chosen across for 3 for the inference. All chains where randomly initialised in the vicinity of the data-generating parameters and the first 2500 iterations were discarded before the ESS computation.

**S9 Appendix. Equivalence of filter inference with a Gaussian filter and ABC based on the mean and variance**. Let *Y*_1_, … *Y*_*N*_ be *N* i.i.d. random, real-valued variables drawn from the data-generating distribution *q*(*y*). Let *q* have nonzero variance, Var[*Y*] *>* 0. Let 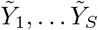 be *S* i.i.d. random variables drawn from the model *p*(*y*|*θ*). Let further 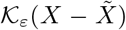 be a kernel with an error margin *ε*, used in ABC to quantify the distance between the data summary statistic *X* = *X*(*Y*_1_,…*Y*_*N*_) and the simulated summary statistic 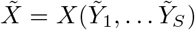. Let the kernel converge to a Dirac delta distribution up to a proportionality factor as the error margin goes to zero, 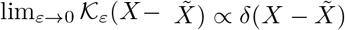. Let further 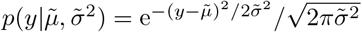 denote a Gaussian filter and 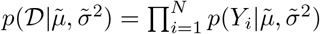 denote its likelihood, where 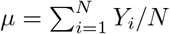 denotes the mean estimator and 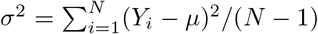 denotes the variance estimator. Then, a Gaussian filter converges to a Dirac delta distribution between (*μ, σ*^2^) and 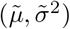 up to a proportionality factor as the number of measurements goes to infinity, 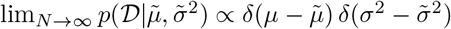. As a result, filter inference with a Gaussian filter is equivalent to ABC based on the mean and variance, 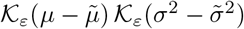, in the limit *N* → ∞ and *ε* → 0.

**Proof:** The maximum likelihood estimators (MLEs) of the Gaussian filter likelihood are

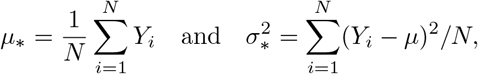

where *μ*_*_ = *μ* and 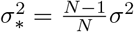. The MLEs are the unique maximum of the Gaussian filter likelihood. The curvature of the log-likelihood at the MLEs is

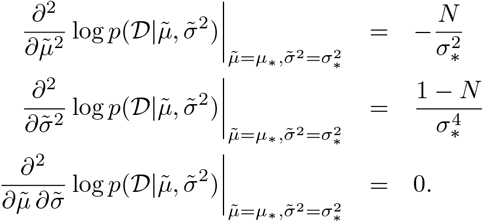

In the limit *N* → ∞, the curvature of the log-likelihood at the MLEs tends to infinity. Due to the monotonicity of the logarithm this implies that the curvature of the likelihood tends to infinity at the MLEs, proving that in the limit *N* → ∞ a Gaussian filter likelihood is proportional to a Dirac delta distribution, 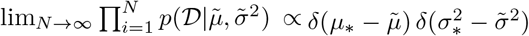. In this limit, the maximum likelihood estimator of the variance coincides with the variance estimator, 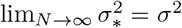, proving the equivalence between filter inference with a Gaussian filter and ABC based on the mean and the variance in the limit *N* → ∞ and *ε* → 0.

**S10 Appendix. Equivalence of filter inference with identifiable summary statistics-based filters and ABC based on the same summary statistics**. Let *Y*_1_, … *Y*_*N*_ be *N* i.i.d. random, real-valued variables drawn from the data-generating distribution *q*(*y*). Let *q* have nonzero variance, Var[*Y*] *>* 0. Let 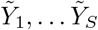 be *S* i.i.d. random, real-valued variables drawn from the model *p*(*y*|*θ*). Let further 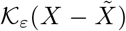 be a kernel with an error margin *ε*, used in ABC to quantify the distance between the data summary statistic *X* = *X*(*Y*_1_, … *Y*_*N*_) and the simulated summary statistic 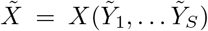. Let the kernel converge to a Dirac delta distribution up to a proportionality factor as the error margin goes to zero, 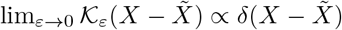. Let further 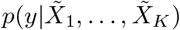 denote a filter defined by *K* summary statistics. Let 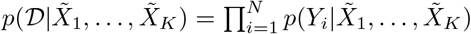 denote its likelihood. Let 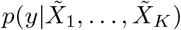 be a *summary statistics-based filter*, when the maximum likelihood estimates of the filter likelihood converge to the summary statistics of the data *X*_1_, …, *X*_*K*_ as *N* → ∞. Let 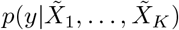 be an *identifiable filter*, when the filter likelihood has a unique maximum, whose curvature tends to infinity in the limit *N* → ∞. Then, an identifiable summary statistics-based filter converges to a Dirac delta distribution between the data summary statistics and the simulated summary statistics up to a proportionality factor as the number of measurements goes to infinity, 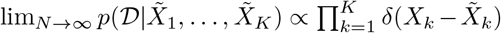. As a result, filter inference with an identifiable summary statistics-based filter is equivalent to ABC based on the same summary statistics, 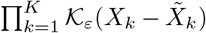 in the limit *N* → ∞ and *ε* → 0.

**Proof:** The MLEs of a summary statistics-based filter recover the summary statistics of the data in the limit *N* → ∞. The MLEs of an identifiable filter are the unique maximum of the filter likelihood, whose curvature tends to infinity as *N* → ∞. As a result, the filter likelihood must be proportional to a Dirac delta distribution at the summary statistics of the data in the limit *N* → ∞, 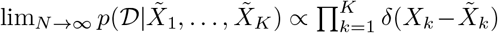. The kernel from ABC also converges to a Dirac delta distribution up to a proportionality factor when the error margin goes to zero, 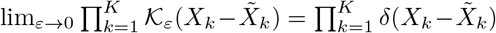. As a result, filter inference with an identifiable summary statistics-based filter is equivalent to ABC based on the same summary statistics in the limit *N* → ∞ and *ε* → 0.

**S1 Table.** Convergence statistics of MCMC chains during the parameter estimation of the early cancer growth model.

|  | 90 IDs (NLME) | 90 IDs | 270 IDs | 810 IDs | 2430 IDs |
| --- | --- | --- | --- | --- | --- |
| $\hat{R}$ of $\mu_{y_0}$ | 1.00 | 1.00 | 1.00 | 1.00 | 1.00 |
| $\hat{R}$ of $\sigma_{y_0}$ | 1.00 | 1.00 | 1.00 | 1.00 | 1.00 |
| $\hat{R}$ of $\mu_\lambda$ | 1.00 | 1.00 | 1.00 | 1.00 | 1.00 |
| $\hat{R}$ of $\sigma_\lambda$ | 1.00 | 1.00 | 1.00 | 1.00 | 1.00 |
| $\hat{R}$ of $\mu_\sigma$ | 1.00 | 1.00 | 1.00 | 1.00 | 1.00 |
| # divergences | 0 | 0 | 0 | 0 | 0 |

**S2 Table. Convergence statistics of MCMC chains during the parameter estimation of the EGF pathway model**.

**S3 Table. Number of log-posterior evaluations**. The table shows the total number of times the log-posterior is evaluated during the inferences in Fig 9.

**S4 Table.** Grid search results: ESS of MH. We ran each MH algorithm with 3 chains for 5000 iterations and computed the ESS after discarding the first 2500 iterations.

| | $\hat{R}$ |
| --- | --- |
| $\mu_p$ | 1.00 |
| $\sigma_p$ | 1.00 |
| $\mu_{k_{\text{on}}}$ | 1.00 |
| $\sigma_{k_{\text{on}}}$ | 1.00 |
| $\mu_{k_{\text{off}}}$ | 1.00 |
| $\mu_{k_{\text{deg},r}}$ | 1.00 |
| $\mu_{k_{\text{deg},a}}$ | 1.00 |
| # divergences | 0 |

|  | 90 IDs | 270 IDs | 810 IDs | 2430 IDs | 2400 IDs (EGF) |
| --- | --- | --- | --- | --- | --- |
| NUTS (warm-up) | 20,000 | 25,626 | 33,059 | 53,589 | 134,413 |
| NUTS (sampling) | 14,985 | 14,985 | 30,969 | 30,969 | 62,937 |
| NUTS (total) | 34,985 | 40,611 | 64,028 | 84,558 | 197,350 |
| Metropolis-Hastings | 90,000 | 90,000 | 90,000 | 90,000 | 180,000 |

| Scale factor | 90 IDs | 270 IDs | 810 IDs | 2430 IDs | 2400 (EGF) |
| --- | --- | --- | --- | --- | --- |
| 1.4 | 71 | **24** | 4 | 3.3 | 3.14 |
| 1.2 | 71 | 15 | 7 | 3.6 | **3.32** |
| 1 | **115** | 12 | **11** | 3.3 | 3.04 |
| 0.8 | 63 | 16 | 6 | 3.2 | 3.22 |
| 0.6 | 82 | 7 | 6 | 3.6 | 3.13 |
| 0.4 | 63 | 8 | 6 | **3.8** | 3.08 |
| 0.2 | 52 | 5 | 5 | 3.4 | 3.29 |

**S5 Table.** Variances of filter posteriors: Early cancer model.

|  | 90 IDs | 270 IDs | 810 IDs | 2430 IDs |
| --- | --- | --- | --- | --- |
| $\text{Var}[\mu_{y_0}]$ | 0.086 | 0.036 | 0.02 | 0.017 |
| $\text{Var}[\sigma_{y_0}]$ | 0.012 | 0.0084 | 0.0046 | 0.0032 |
| $\text{Var}[\mu_\lambda]$ | 0.072 | 0.036 | 0.018 | 0.0014 |
| $\text{Var}[\sigma_\lambda]$ | 0.0082 | 0.00456 | 0.0033 | 0.0024 |
| $\text{Var}[\mu_\sigma]$ | 0.0089 | 0.1176 | 0.0082 | 0.0096 |

**S5 Table.** Variances of filter posterior: EGF pathway model.

|  | Variance |
| --- | --- |
| $\mu_p$ | 0.00005 |
| $\sigma_p$ | 0.0001 |
| $\mu_{k_{\text{on}}}$ | 0.00006 |
| $\sigma_{k_{\text{on}}}$ | 0.00004 |
| $\mu_{k_{\text{deg},r}}$ | 0.0000008 |
| $\mu_{k_{\text{deg},a}}$ | 0.000002 |

**S1 Fig.**
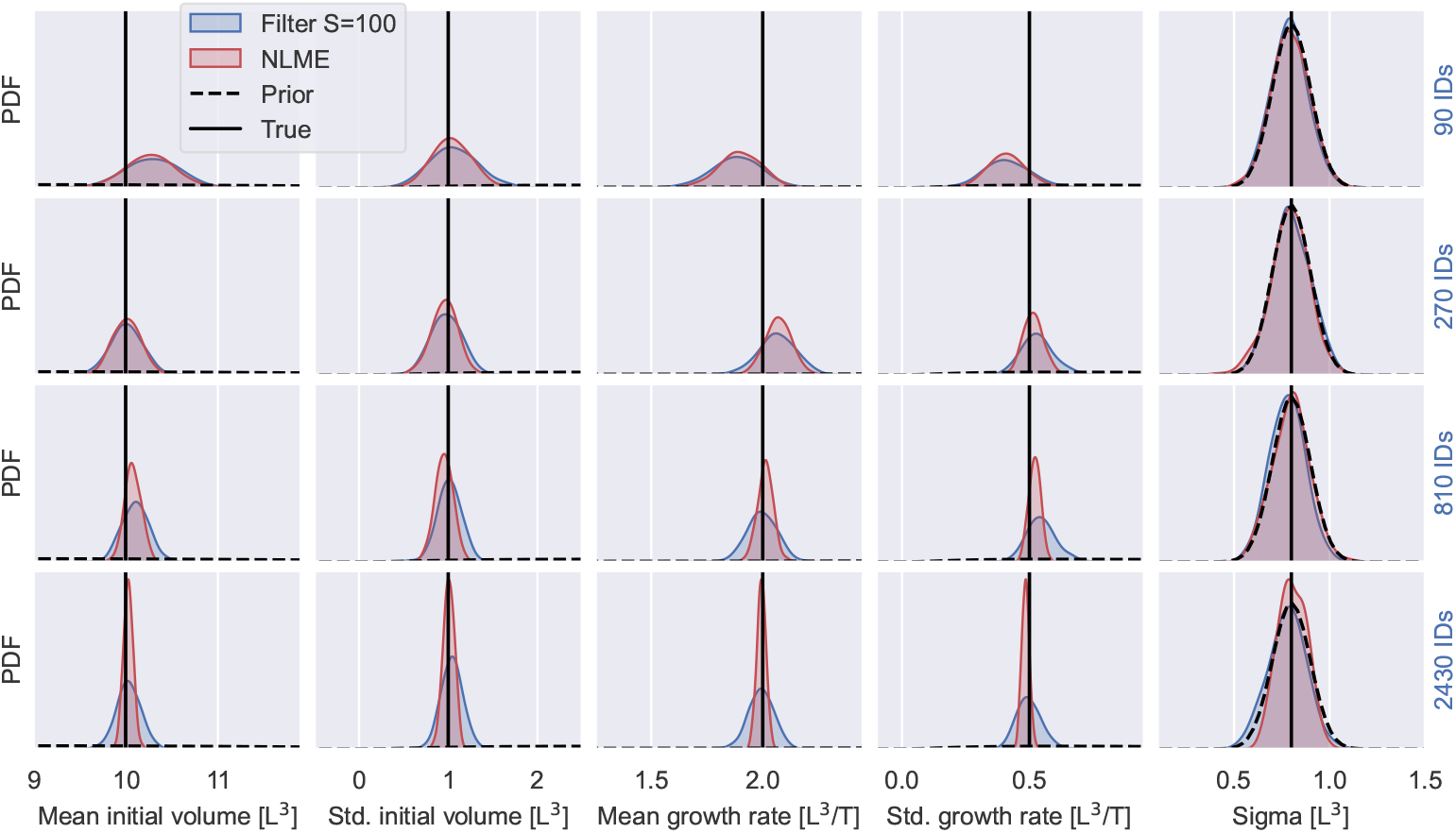
IIV-noise identifiability of early cancer growth model I. The figure shows inference results of the early cancer model using NLME inference and filter inference with a Gaussian filter and *S* = 100 simulated individuals. The parameters are estimated from snapshot datasets with measurements from *N* = 90, *N* = 270, *N* = 810, *N* = 2430 individuals. The measurements are generated for 6 time points between 0 and 0.6 as described in Early cancer growth model inference. The first row shows the posterior distributions inferred from measurements of *N* = 90 individuals. The second, third and fourth row analogously show the results for *N* = 270, *N* = 810 and *N* = 2430. The filter posteriors are illustrated in blue and the NLME posteriors are illustred in red. The prior distributions are illustrated by black dashed lines and the data-generating parameter values are depicted by solid black lines. *θ* = (10, 0.7, 2, 0.5, 1.127), is illustrated in red.

**S2 Fig.**
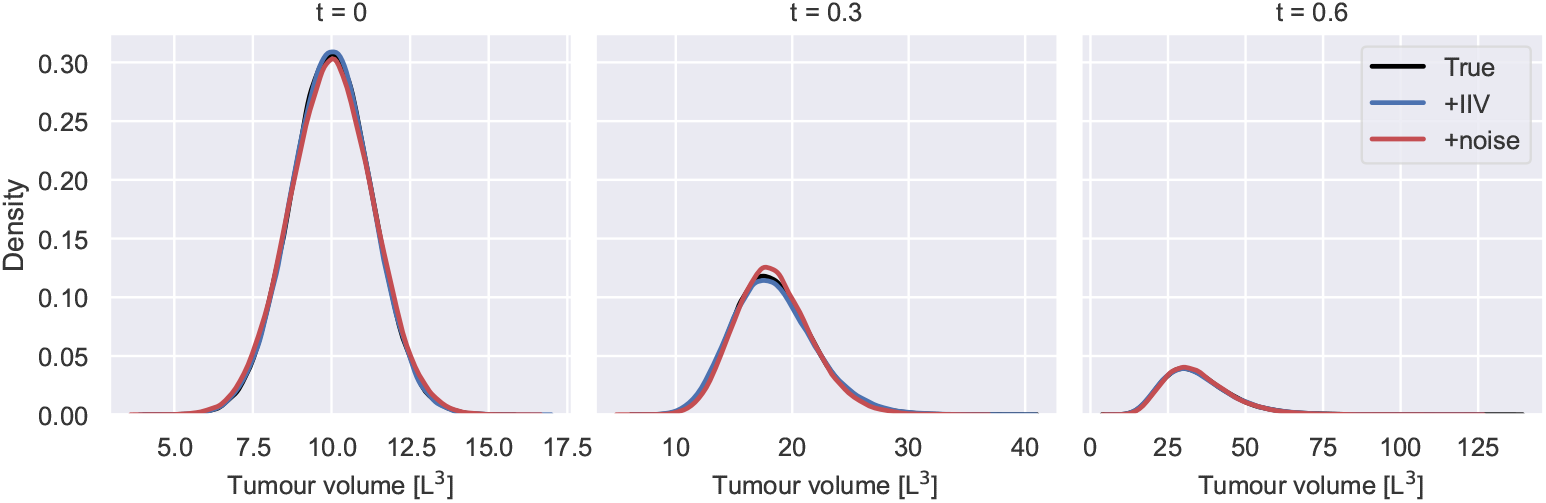
Early cancer growth measurement distribution. The figure is an extension of S1 Fig and shows the measurement distribution *p*(*y*|*θ, t*) of the early cancer growth model at *t* = 0, *t* = 0.3 and *t* = 0.6 for three different sets of parameter values. The measurement distribution corresponding to the data-generating parameters, 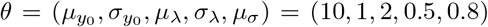, is shown in black. A measurement distribution that overestimates the IIV contributions to the measurement variability, *θ* = (10, 1.195, 2, 0.5, 0.47), is illustrated in blue, and a measurement distribution that overestimates the noise contributions to the measurement variability, *θ* = (10, 0.7, 2, 0.5, 1.127), is illustrated in red.

**S3 Fig.**
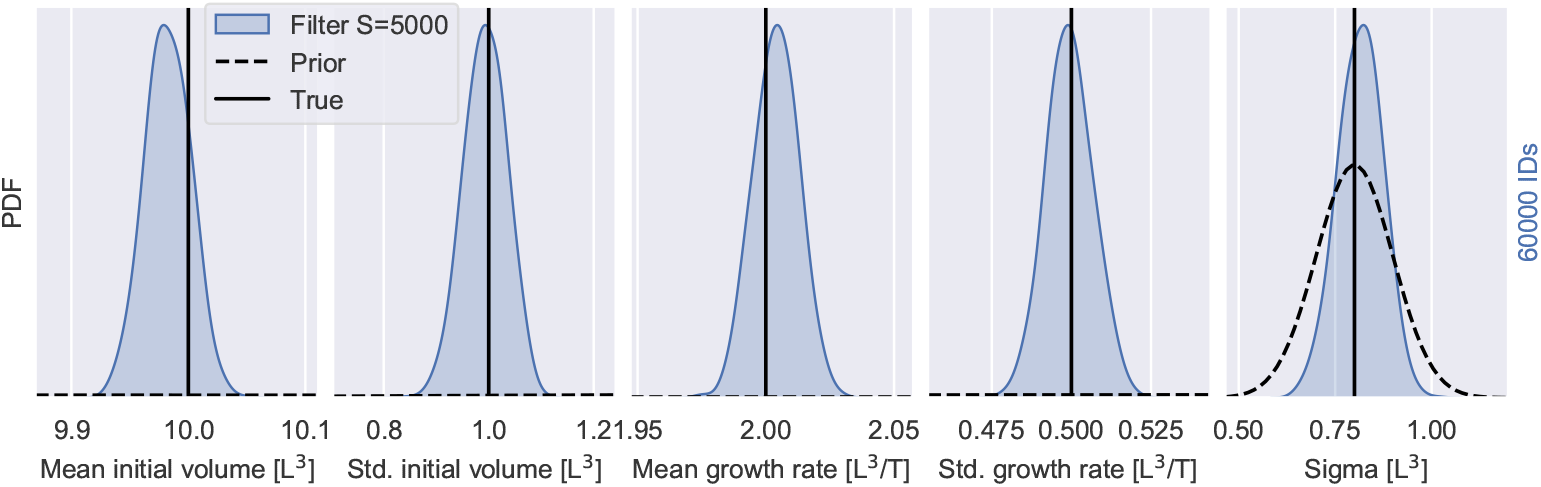
IIV-noise identifiability of early cancer growth model II. The figure is an extension of S1 Fig and shows filter inference results for the early cancer growth model from 60 000 snapshot measurements using Gaussian filters with *S* = 5 000 simulated individuals. The prior distributions are illustrated by black dashed lines and the data-generating parameter values are depicted by solid black lines.

**S4 Fig.**
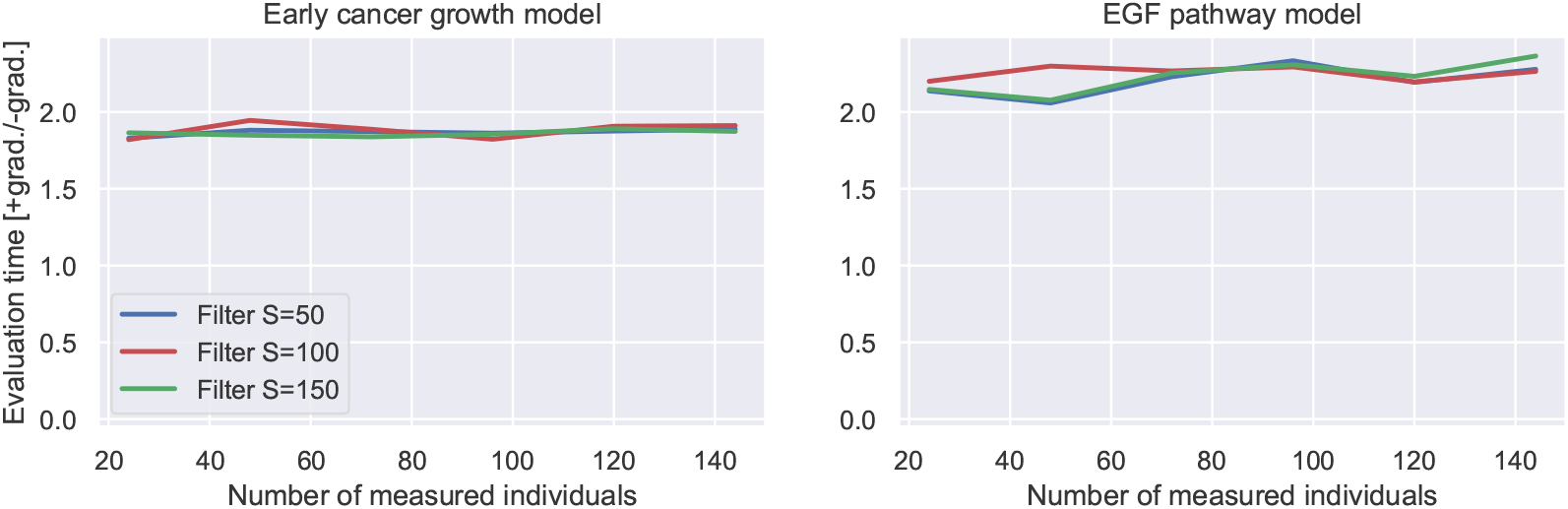
Computational costs of log-posterior evaluation with and without gradients. The figure is an extension of Fig 7 and shows the evaluation time of the filter log-posterior with gradients in units of the evaluation time of the filter log-posterior without gradients for different numbers of measured individuals. The evaluation times are estimated according to S7 Appendix. The left panel shows the results for the early cancer growth model and the right panel the results for the EGF pathway model for *S* = 50 (blue), *S* = 100 (red) and *S* = 150 (green) simulated individuals.

## Notes

https://github.com/DavAug/filter-inference

### Competing Interest Statement

KW and ACW are employees and shareholders of F. Hoffmann-La Roche Ltd.

